# Muscle spatial transcriptomic reveals heterogeneous profiles in untreated juvenile dermatomyositis and the persistence of pathological signature after remission

**DOI:** 10.1101/2025.04.15.648883

**Authors:** Margot Tragin, Séverine A. Degrelle, Baptiste Periou, Benjamin Chaigne, Brigitte Bader-Meunier, Christine Barnerias, Christine Bodemer, Isabelle Desguerre, Mathieu Paul Rodero, François Jérôme Authier, Cyril Gitiaux

**Author notes:** **Corresponding Author:** Pr Cyril Gitiaux. Hôpital Necker-Enfants Malades, 149 rue de Sèvres, 75015 Paris, France. these senior authors contributed equally to the work (Mathieu Paul Rodero, François Jérôme Authier, Cyril Gitiaux).

## Abstract

This study aimed to investigate the spatial heterogeneity of molecular signature in the muscle of juvenile dermatomyositis (JDM) patients before and after treatment in comparison to healthy paediatric muscle tissue. Unsupervised reference-free deconvolution of spatial transcriptomics and standardized morphometry were performed in two JDM muscle biopsies with different clinical severity at disease onset and compared to healthy paediatric muscle. In a second step, identified signatures were scored in two additional JDM muscle biopsies from the same patient before and after remission. Disappearance of the normal muscle signature mostly corresponding to mitochondrial biology was observed in JDM. Three pathological transcriptomic signatures were isolated, related to “myofibrillar stress”, “muscle remodeling” and “interferon signaling” signatures. The “myofibrillar stress signature” was prominent in the most severe biopsy while the “muscle remodeling” signature was mostly present in the biopsy from the patient with good outcome. These signatures unveiled genes not previously associated with JDM including *ANKRD1* and *FSLT1* for “myofibrillar stress” and “muscle remodeling” signatures, respectively. Post-treatment analysis of muscle after two years of remission showed a persistence of pathological signatures. This study of JDM muscle identified spatially distributed pathological signatures that persist after remission. This work paves the way for a better understanding of the pathophysiology in affected muscle and the identification of biomarkers that predict relapse.

## INTRODUCTION

Juvenile dermatomyositis (JDM) is a rare and heterogeneous autoimmune and inflammatory myopathy characterized by specific skin and muscle involvement. Other organ involvements, particularly of the lungs and gastrointestinal tract, are associated with mortality in some cases. Although modern treatment has improved the outcome of JDM, cohort studies suggest that only 50% of patients achieve remission off therapy [1, 2]. Muscle-specific autoantibodies (MSA) help to stratify patients into different phenotypes with implications for prognosis and responses to treatment [3]. JDM pathophysiology is characterized by a strong and selective upregulation of type I interferon (IFN-I, i.e IFNα/β) in the circulation and tissue which is attributed to complex interplay between genetic and environmental risk factors [4, 5]. In muscle tissue, the preferential activation of IFN-I pathway is a major contributor to myopathological changes and is associated with a multifocal capillary loss, muscle ischaemia and perifascicular myofibre atrophy. Altered mitochondrial biology also occurs in JDM and may be part of the pathological loop that drives IFN-I production [6]. Deregulation of IFN-I pathway is interconnected with endothelial dysfunction and vasculopathy and is associated with the poorer prognosis of JDM [7]. The nature of the triggering events and the link between microvascular dysfunction and IFN-I upregulation remains unresolved [8, 9]. Early identification of patients with more severe diseases requiring specific treatment remains a major unmet medical need. Such identification remains hampered by the poor understanding of the cellular and molecular mechanisms underlying the variability of disease progression. Here, we took advantage of multi-cellular pixel resolution spatial transcriptomic (ST) profiling of muscle using a reference-free deconvolution approach in JDM to describe spatial transcriptomic signatures at disease onset. Furthermore, ST showed that abnormal signatures persist after remission, paving the way for understanding of disease relapse in JDM.

## MATERIALS AND METHODS

### Muscle biopsies

Patients were recruited in the reference centers for Rare Pediatric Inflammatory Diseases (RAISE/FAI2R) and for Rare Neuromuscular Diseases (Nord-Est-Ile de France/FILNEMUS). JDM patients were classified according to conventional clinico-pathological criteria [10]. Deltoid biopsies from JDM patients were performed at the time of the diagnosis in the context of diagnostic work-up. Control biopsies were obtained from the Henri Mondor hospital muscle biobank. All biopsy samples were blindly reviewed for clinical data using the severity scoring tool for muscle biopsy evaluation in JDM patients [11].

### Methods for histology, immunohistochemistry and morphometric analysis: see supplementary information

#### Visium spatial Gene expression library construction

Spatial gene expression libraries were generated following the 10X Genomics Visium Spatial Gene Expression protocol (User Guide, CG000239 Rev F). Muscle 10 µm-cryo-sectionned were placed onto Visium glass slides. The slides were transferred to 37°C for 1 min before immersion and fixation in methanol at -20°C for 30 min. After fixation, sections were dried by adding 500 μl isopropanol for 1 min then air-dried, stained with Mayer’s Hematoxylin/Eosin and dried again for 5 min at 37°C. Imaging was performed using a Zeiss Axio Imager.D1/D2 at 10x magnification, and the images were processed using ZEISS ZEN 3.7 software. After imaging, the slide was transferred to the Slide Cassette. The tissue sections then were permeabilized using the permeabilization enzyme at 37°C (incubation time was determined following tissue permeabilization optimization), after which it was removed and each well washed with 0.1x SSC. A reverse transcription mix containing RT Reagent, Template Switch Oligo, Reducing Agent B, RT Enzyme D, and Nuclease-free water were added, and the slide was incubated at 53°C for 45 min. After removal, 0.08M KOH was added to each well and incubated at room temperature (RT) for 5 min, then washed with EB (Qiagen, 19086). Next, a second strand mix containing Second Strand Reagent, Second Strand Primer, and Second Strand Enzyme was added and incubated at 65°C for 15 min. After washing with EB, 0.08M KOH was added to each well and incubated for 10 min at RT. Into each tube of an 8-tube strip, 5 μl Tris (1M, pH7.0) was added, followed by 35 μl of sample from the wells. cDNA amplification was carried out using a master mix containing Amp mix and cDNA primers. The number of cycles was determined using qPCR with the addition of KAPA SYBR FAST (Sigma-Aldrich, KK4600). The samples were purified using SPRIselect (Beckman Coulter, B23318) at 0.6X. Quality control was performed on all cDNA samples using an Agilent TapeStation Screen Tape, and concentrations were determined using a High Sensitivity Qubit assay (Thermo-Fisher). Next, fragmentation was carried out by incubating the samples at 32°C for 5 min and then 65°C for 30 min with EB and a fragmentation mix containing Fragmentation Buffer and Fragmentation Enzyme. After fragmentation, a purification using SPRIselect at 0.6X and 0.8X was performed before adaptor ligation. An adaptor ligation mix containing Ligation Buffer, DNA Ligase, and Adaptor Oligos were added and incubated for 15 min at 20°C, followed by purification using SPRIselect at 0.8X. For indexing, Amp mix, and an individual dual index (Kit TT Set A) were added to each sample, and the indexing protocol was run in a thermocycler. The number of cycles for indexing was determined by the cDNA input calculated from the previous quality control. After indexing, a final purification was performed using SPRIselect at 0.6X and 0.8X. Quality control was performed on all samples using an Agilent TapeStation Screen Tape, and concentrations were determined using a High Sensitivity Qubit assay (Thermo-Fisher). Finally, the libraries were sequenced in paired-end 150 bp mode on an Illumina NextSeq 500 system. The sequencing depth was determined by the percentage of capture areas covered by each muscle sample.

#### Visium data processing and analysis

Illumina reads of each area were aligned on the GRCh38-2020-A (Ensembl 98) reference and counted using the spaceranger count function from the 10X genomics software Spaceranger 2.1.1. Areas corresponding to the same experiment were integrated using spacerange aggregate function with default parameters. Exploration of integrated data using the 10X Genomics software Loupe Browser 7.0.1 allows to annotate and discard spots outside and/or associated with low quality areas of the biopsies (e.g. edges, folded slice…). All further analyses and visualization were performed under R (version 4.4.2). Reference-free deconvolution was performed on the count matrix using the STdeconvolve v1.4.0 R package [12]. Spots with a minimum of 100 detected genes and informative genes reaching the restrictCorpus default parameters were considered. The number of signatures was fixed a priori to K=5, corresponding to a local minimum of perplexity **(online supplemental Figure S1)**. Then, a list of marker genes (up and downregulated) was defined in each of the 5 signatures based on the following log2FC formula [12]. Enrichment tests using ReactomePA v 1.46.0 R package [13] and Ingenuity Pathway Analysis (IPA, Qiagen Redwood City), were performed on marker genes list. For IPA, pathway network with significant p-values (p < 0.05) was generated [14]. In a next step, the correspondence of spatial transcriptomics data from JDM patient biopsies sampled before and after treatment with the transcriptomics signatures (i.e. list of marker genes) were investigated using a directional (upSet and downSet options) Singscore v. 1.20.0 R package [15, 16]. Singscore ranges from -1 to 1, a spot which singscore is 1 means that its transcriptome matches the tested signature. Empirical Pvalue were estimated for each scored spot using 1000 permutations with generateNull() and getPvals() functions from Singscore v. 1.20.0 R package. Estimated Pvalue corresponds to the results of an enrichment test of the signatures in each spot. Dimensional reduction and t-SNE (t-distributed stochastic neighbor embedding) calculation were performed using M3C v.1.22.0 R package [17]. Data visualization was done with ggplot2 v. 3.5.0 R package.

## RESULTS

### Spatial transcriptomic analysis reveals specific and spatialized muscle signatures in JDM as compared to healthy muscle

To define the transcriptomic signature of the different areas of affected muscle in JDM, we performed 10X Genomics Visium Spatial Gene Expression analysis on the muscle obtained at diagnosis of an acute NXP2+ JDM with severe muscle involvement (muscle biopsy severity score: 20/27, VAS: 7/10), an indolent, slowly progressive MSA-negative JDM (muscle biopsy severity score: 13/27, VAS: 4/10), and on one healthy muscle (HM). The clinical and demographic characteristics of the JDM patients are shown **in Table 1**. Morphometric analysis of the myofibre size (cross-sectional area), of fibrosis and capillary density showed a more severe overall atrophy in indolent JDM whereas fibrosis and capillary loss were similar in acute and indolent JDM as compared to the normal muscle tissue **(Figure 1A-L)**. On average, 80 million RNA sequences were produced per biopsy ([44 x 10^6^;135 x 10^6^], SD: 48 x 10^6^), covering a total of 2071 usable spots. The capture of 8,800 to 90,600 (SD 17,750) reads per spot allowed the detection of a mean of 336 ([252; 398] SD: 75.4) genes per spot, corresponding to approximately 14 000 genes detected per entire biopsy. The mean number of genes per spot was consistent across the 3 biopsies (acute: 398, indolent: 358, HC: 252). T-SNE performed on global aggregated data showed that spots from both muscles of JDM patients clustered together, away from the healthy muscle spots **(Figure 2A)**. In accordance with the clinical and pathological severity, the distribution of spots in the t-SNE shows a gradient, with spots from the indolent JDM muscle mostly positioned between spots from the healthy muscle and spots from the acute JDM muscle.The reference-free deconvolution approach allowed i) to identify 5 transcriptional profiles (i.e. molecular signatures) **(online supplemental Figure S1)**, in which a list of specifically up or down-regulated transcripts were identified **(online supplemental Table S1)** and ii) to estimate the percentage of each molecular signature in each spot of each biopsy. Signature 1 was prominent in normal muscle (99,3%), signature 2 was less expressed and similar between the three conditions, signature 3 was prominent in acute JDM (78,8%) while signature 5 was prominent in indolent JDM muscle (65,3%). Signature 4 was associated with both JDM (acute JDM 9,2%; indolent JDM 14,2%) **(Figure 2 B-C)**. This analysis refined the transcriptomic mapping on the 3 biopsies and showed that the transcriptome of JDM patients is spatially distributed, with regionalized specific signatures, whereas healthy biopsy showed a homogeneous transcriptome in each spot. These transcriptomic profiles reflected the respective histopathological patterns observed in the two JDM muscles **(online supplemental Figure S2)**, namely ischemic injuries in acute JDM with microinfarcts, myosinolysis and perifascicular atrophy, and chronic moderate myopathic changes in indolent, slowly progressive JDM with myofiber size variation and internalized nuclei.

**Table 1.** Demographics and clinical characteristics of JDM patients included in the study.

| Sample ID | JDM1 | JDM2 | JDM3 onset | JDM3 treated |
| --- | --- | --- | --- | --- |
| Age at sample, years | 13 | 5 | 13 | 15 |
| Sex | F | F | F | F |
| Disease duration at time of biopsy, months | 1 | 4 | 1 | 24 |
| Myositis specific antibody | NXP2 | 0 | NXP2 | NXP2 |
| Disease activity at time of biopsy: |  |  |  |  |
| CMAS (0-52) | 2 | 30/46 | 2 | 52 |
| MMT (0-80) | 47 | NR | 38 | 80 |
| CK U/L | 3020 | 1500 | 862 | 100 |
| Characteristic skin rash | 1 | 1 | 1 | 0 |
| Gottron's papules | 1 | 1 | 1 | 0 |
| Telangiectasiae | 1 | 0 | 1 | 0 |
| Subcutaneous limb edema | 1 | 0 | 1 | 0 |
| Skin ulcerations | 0 | 0 | 0 | 0 |
| Interstitial lung disease | 0 | 0 | 0 | 0 |
| Cardiovascular involvement | 0 | 0 | 1 | 0 |
| Gastrointestinal involvement | 0 | 0 | 1 | 0 |
| Muscle biopsy score data* : |  |  |  |  |
| Inflammatory domain (0-12) | 8 | 4 | 12 | 0 |
| Muscle fibre domain (0-10) | 7 | 7 | 9 | 2 |
| Vascular domain (0-3) | 3 | 1 | 3 | 0 |
| Connective tissue domain (0-2) | 2 | 1 | 2 | 1 |
| Total biopsy score (0-27) | 20 | 13 | 26 | 3 |
| Histopathology visual analogue score (0-10) | 7 | 4 | 8 | 1 |

\*Data generated using validated JDM muscle biopsy score from Varsani et al [11]

**Figure 1.**
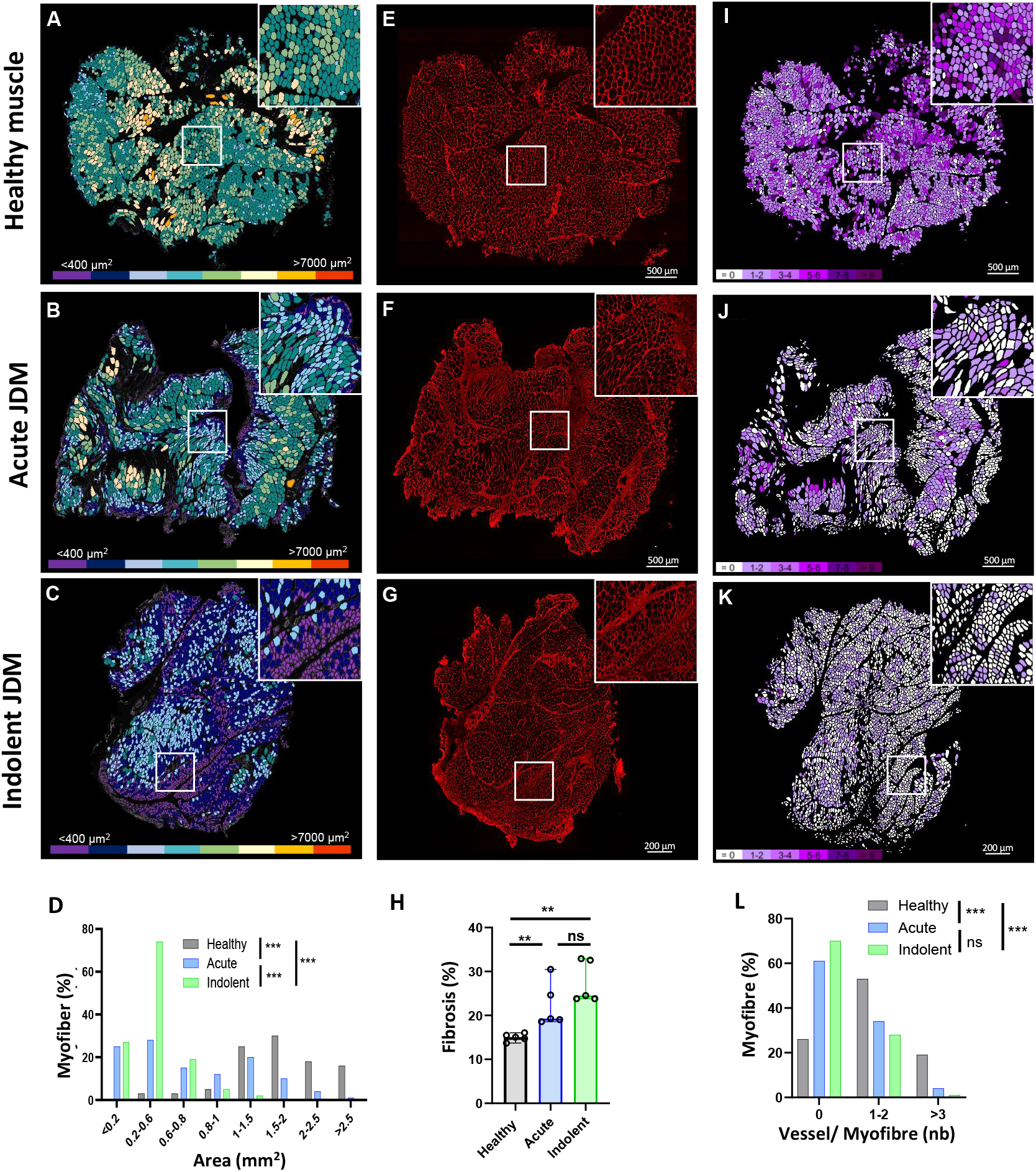
Detailed automated morphometric analysis of the muscle biopsies used to determine spatial transcriptomic signatures in JDM. Immunofluorescence staining of healthy muscle **(upper panel: A-E-I)**, acute JDM **(middle panel: B-F-J)** and indolent JDM **(lower panel: C-G-K)**. Representative images processed by an ImageJ macro **(A-B-C)** of myofibres areas (from ≤ 400 to > 7000 μm^2^) performed using a muscle tissue section stained for laminin to delineate myofibres; **(E-F-G)** of collagen VI immunofluorescence staining as marker of fibrosis; **(I-J-K)** of vessel number (CD31+) per myofibre (from 0 to > 9) performed using a muscle tissue section double-stained for laminin and CD31. Tool for the ImageJ software allowing for automated morphometry of **(D)** myofibre cross-sectional areas **(H)**, of fibrosis and **(L)** endomysial vascular density assessed by the number of capillary associated with each myofibre. Bar graphs: median with range shown. Statistical analysis: chi-square **(D, L)**; Kruskall Wallis allowed by Dunn’s multiple comparison test **(H)**. JDM, juvenile dermatomyositis. NS: non-significant. p values * ≤0.05 ** ≤ 0.01 *** ≤ 0.01.

**Figure 2.**
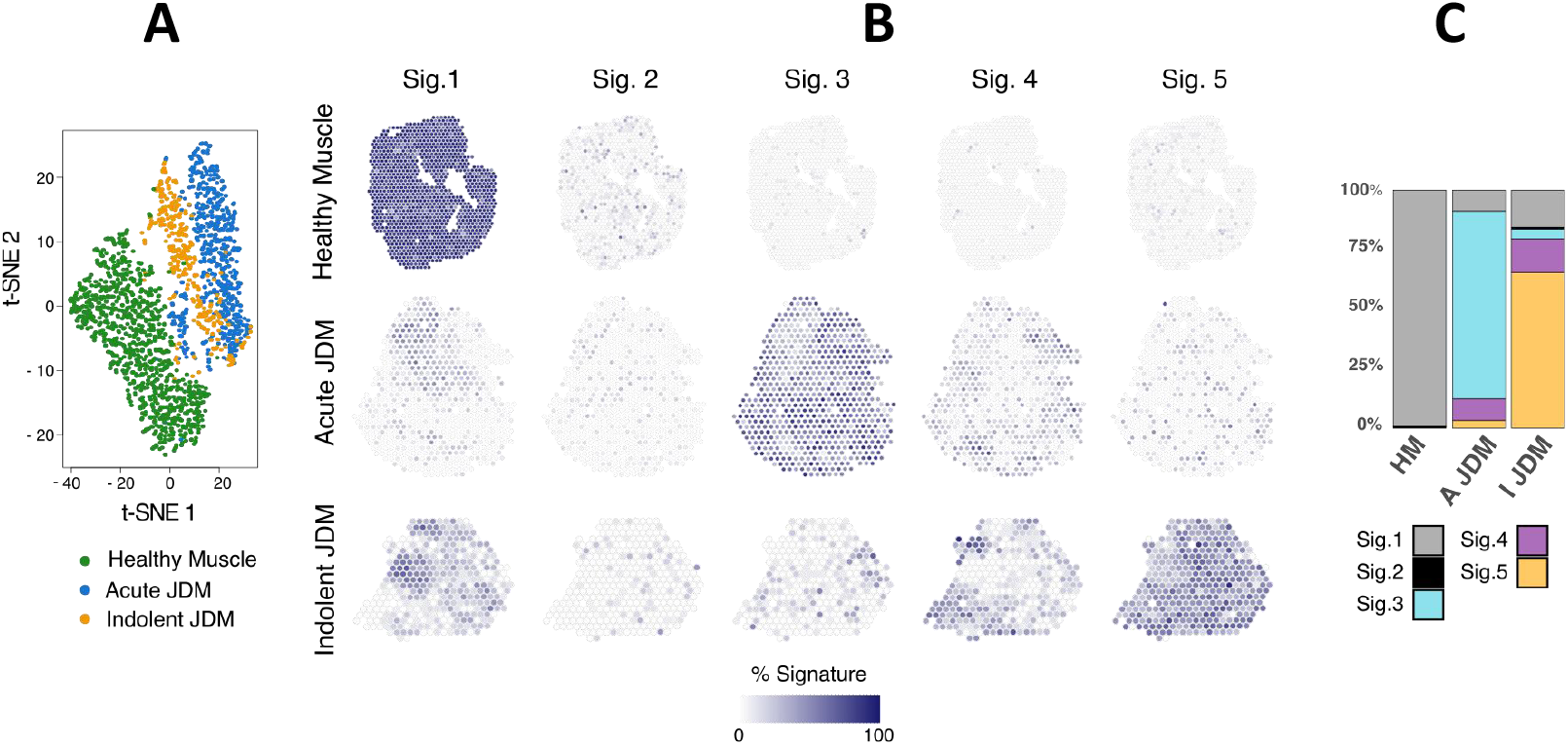
Spatial transcriptomic analysis reveals specific and spatialized muscle signatures in JDM as compared to healthy muscle. T-distributed stochastic neighbor embedding (t-SNE) dimensional reduction of 2071 transcriptomes (spots) of the 3 biopsies **(A)**. *De novo* deconvolution of transcriptomes (spots) in 5 signatures. Healthy muscle (first row), acute JDM (middle row) and indolent JDM (last row). Signature 1 is mostly associated with healthy muscle, signature 3 with acute JDM and signature 5 with indolent JDM. Signature 4 is present in both JDM and signature 2 is not specific and present among the 3 conditions **(B)**. Bar graph of the percentages of each signature in the three conditions **(C)**.

### Muscles from acute and indolent JDM could be distinguished by the expression of a predominant myofibrillar stress signature or an extracellular matrix remodeling signature respectively but shared IFN signaling signature

Using the Reactome database, we identified significantly enriched pathway for each of the 5 lists of genes previously identified **(Figure 3ACEG, online supplementary Table S2)**. Signature 1, which was mostly present in healthy muscle, was mainly characterized by higher expression of genes related to “citric acid (TCA) cycle and respiratory electron transport”. Indeed, *ATP6, ATP8* and *NDA5* were among the most highly expressed genes in healthy muscle compared to JDM. Signature 3, which was mostly present in the acute JDM, was associated with the “muscle contraction” pathway, with *ACTC1, MYH8* and *MHY3* among the top over expressed genes compared to the other biopsies. Signature 5, which is mostly present in the indolent biopsy was associated with the “Extracellular matrix organization” pathway. The top overexpressed genes from this signature are associated with extracellular matrix composition (exp *COL6A3/COL15A1*), with mitochondrial biology (exp *NDUFB5/NDUFB9/NDUFA7*) and with other genes of interest (exp *FSTL1)*. Interestingly, signature 4 which was common between the 2 JDM biopsies was strongly associated with the “interferon signaling” pathway, notably with *ISG15, IFIT1* and *IFI27* among the most overrepresented transcripts. Signature 2 was almost absent in all biopsies (less than 1 %). It was characterized by the overexpression of genes shared with signature 1 **(online supplemental Figure S3 and online supplemental Table S2)**. For this reason, we decided not to include it for the rest of the analysis. Interestingly, network analysis performed with IPA showed very comparable results for Signature 1, 3 and 4 **(Figure 3BDFH)**. However, the most abundant network for signature 5 is composed of genes from the mitochondrial respiratory chain (*NDUFB9, B5* and *A7*) as well as genes of interest not previously reported in muscle from JDM (for exp *FSTL1* and *TAGLN2*) confirming the complexity of this signature 5 likely corresponding to muscle remodeling **(Figure 3H)**. Based on these results, we relabeled signature 1 as “healthy muscle signature”, signature 3 as “myofibrillar stress signature”, signature 4 as “interferon signature” and signature 5 as “muscle remodeling signature”.

**Figure 3.**
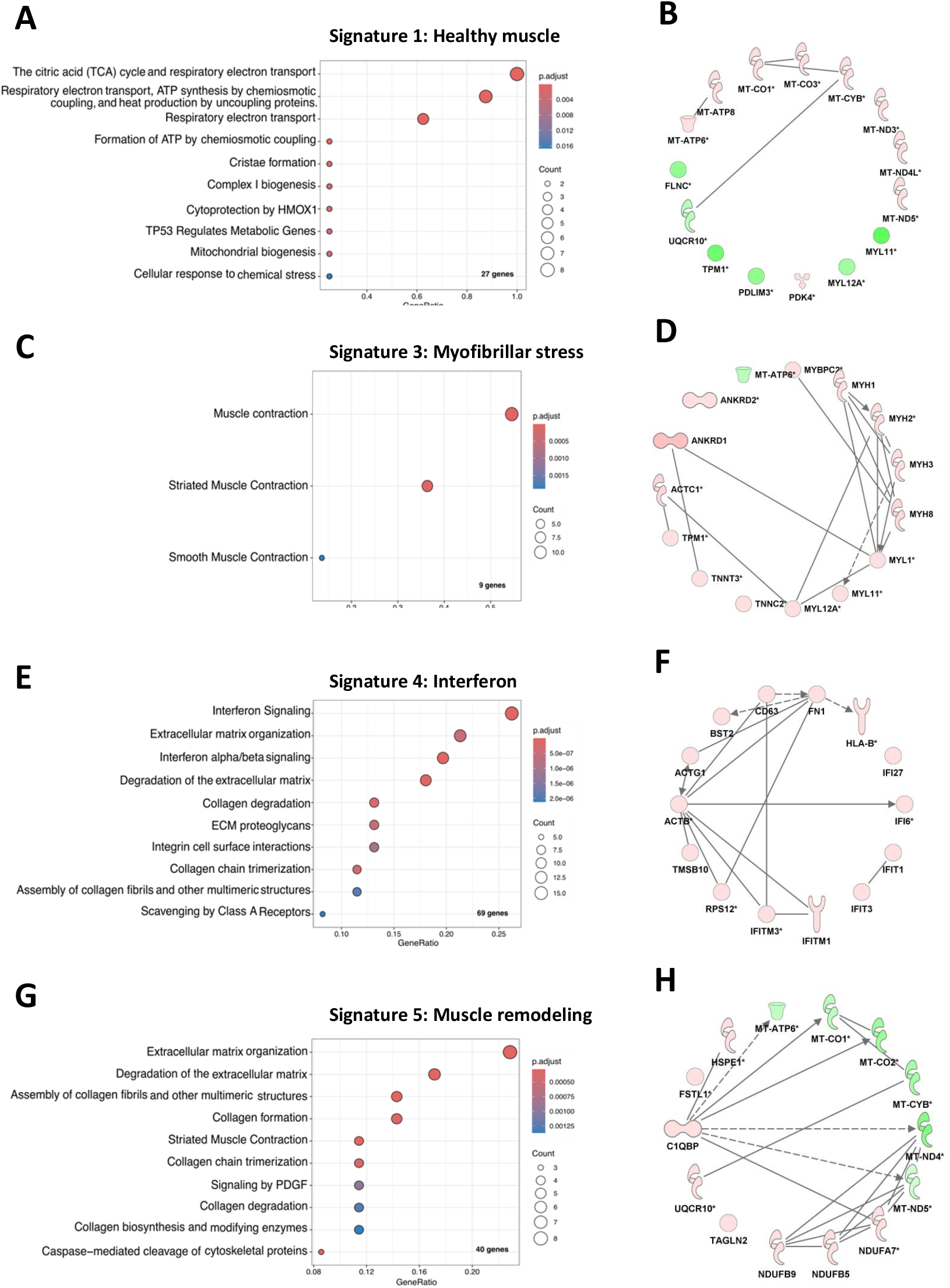
Indolent and acute muscles display distinct transcriptomic signatures. Enriched Reactome pathways **(A-C-E-G)** and IPA analysis **(B-D-F-H)** were performed on differentially expressed genes from signatures 1, 3, 4 and 5 respectively. Complete list of pathways and results for signature 2 are available in supplemental figure S2. For Reactome pathways analysis, only the ten most significant pathways are shown (com. The count represents the number of genes from marker genes list, that are found in a specific Gene Ontology (GO) term. The GeneRatio is the proportion of marker genes associated with a particular GO term relative to the total number of our marker genes in the lists. For IPA analysis, genes up- and down-regulated are respectively highlighted in green and red. The different shapes represent various types of molecules and their relationships according to the IPA knowledge base (http://ingenuity.com/).

### Persistence of myofibrillar stress signature and muscle remodeling signature in the muscle of a patient in two-years remission after JAK inhibitor treatment

We then investigated the correspondence of spatial transcriptomic data from two additional muscle biopsies obtained at diagnosis and after two years of remission from the same JDM NXP2+ patient as compared to healthy muscle with the transcriptomic signatures identified above **(Figure 4A-C)**. At disease onset the patient presented with a very severe ischemic myopathy characterized by massive endomysial capillary loss (muscle severity score: 26/27, VAS:8/10). The second muscle biopsy performed 42 months later and after 22 months of JAKi treatment showed almost no abnormality: Moderate residual changes (myofibre size irregularity, rounded appearance and centronucleation of myofibres), mild endomysial fibrosis and complete restoration of endomysial microvascular bed (muscle severity score: 3/27, VAS:1/10). To quantify the change in signature expression, we calculated a Singscore for each spot of each biopsy representing the relative expression of each signature within this spot. Compared to healthy muscle, the “healthy muscle” signature was almost absent in biopsy performed at diagnosis, while all three pathological signatures “myofibrillar stress”, “interferon” and “muscle remodeling” were present. Following JAKi treatment, we observed a significant increase in the healthy signature in most spots. Conversely, we observed a decrease in all three pathological signatures. Interestingly, only the interferon score returned to levels comparable to healthy muscle, but unexpectedly both the “myofibrillar stress” and “muscle remodeling” signature persisted in the biopsy **(Figure 4D)**.

**Figure 4.**
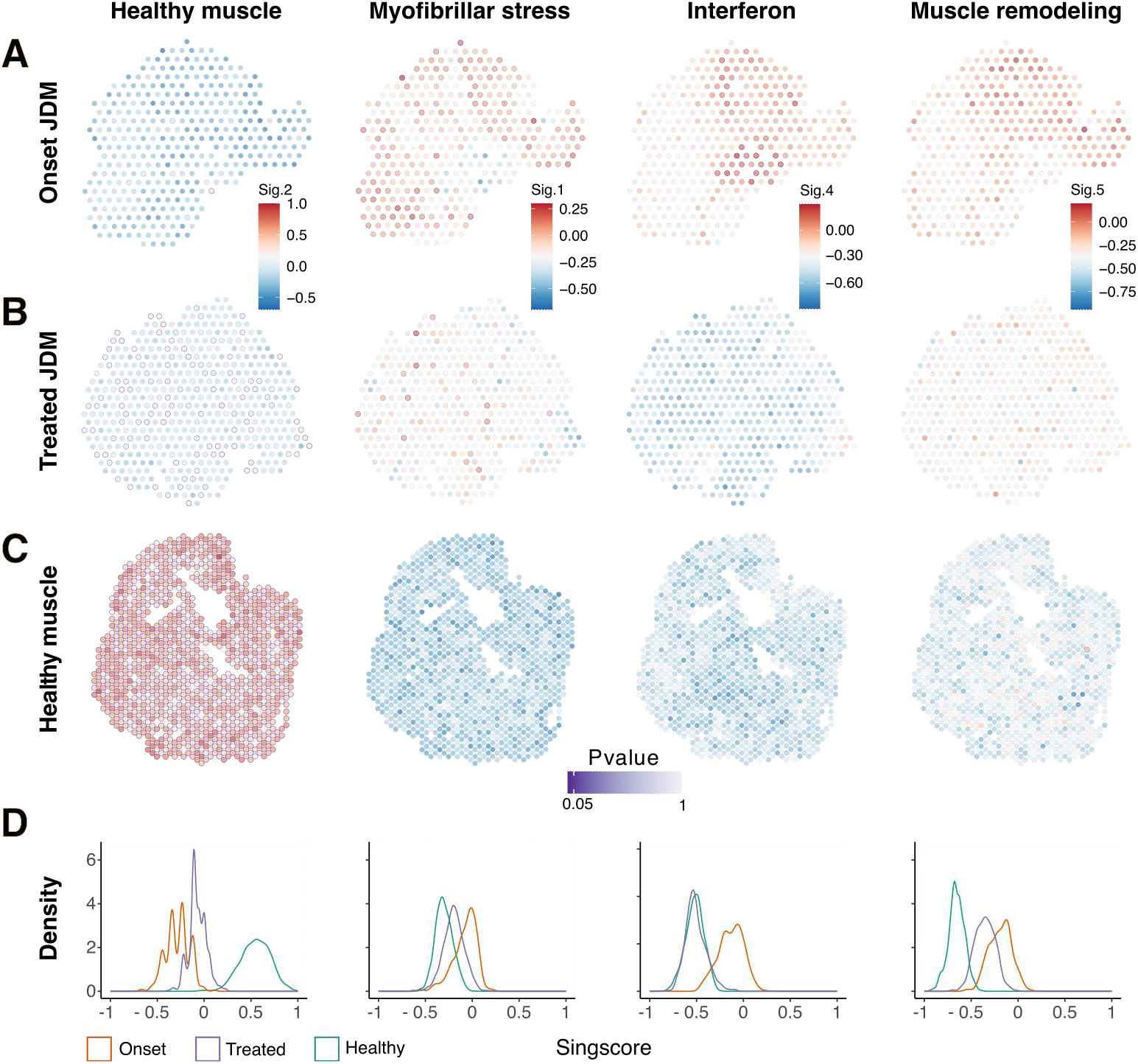
Persistence of myofibrillar stress signature and extra cellular matrix remodeling signature in the muscle of a patient in two-years remission after JAK inhibitor treatment. Directionnal singscore and associated Pvalue (purples color scale) calculated by permutation tests of marker genes lists for lists for the 4 normal and pathological muscle spatial transcriptomic signatures (healthy, myofibrillar stress, IFN signaling and muscle remodeling): Singscores in each spatial transcriptomic spot: disease onset JDM biopsy **(A)**, after JAKi treatment JDM biopsy from the same patient **(B)**, healthy muscle **(C)**. Distribution of singscores values (from -1 to 1) in the 3 muscle biopsies **(D)**.

## DISCUSSION

Our understanding of the pathophysiology of JDM is based on studies conducted over the past decades, initially focusing on muscle vasculopathy and more recently on the role of IFN-I upregulation. The lack of significant progress in characterizing the drivers of disease onset and progression has limited the identification of sensitive biomarkers of treatment response and the development of new targeted therapies. In particular, we have little insight into the chronology of events taking place in the affected muscle. To date, omics studies in DM/JDM have focused on either circulating cells or bulk muscle and single cell/nuclei RNA sequencing. These studies have identified distinct gene expression patterns in myositis associated with different autoantibodies [18, 19] questioning the pathological significance of these autoantibodies [20] or have addressed the mechanisms contributing to the abnormal IFN-I signature in the patients [6, 21].

The recent development of spatial transcriptomics constitutes a breakthrough in the understanding of pathological mechanisms at the tissue level and has recently been applied to several neuromuscular disorders in animal models [22, 23] and in human myositis, including sarcoidosis myositis [24], inclusion body myositis [25] and dermatomyositis [26]. The latter study showed mitochondrial damage gene signature in myofibres, which was accentuated when located in contact with immune cell infiltrates supporting a crosstalk between inflammatory cells and muscle mitochondria in myositis [26]. Here, for the first time, we have applied this strategy to muscle biopsies from juvenile dermatomyositis. Visium technology allows the investigation of 55 μm circular spots (i.e. subcellular resolution), in which mRNA from several biological cell types could be captured [27]. There are two approaches to investigate the diversity of cell types in each spot: i) deconvolution using a reference [28] consists in identifying biological cell types based on a single cell/nuclei reference database and ii) reference-free deconvolution [12], which allows the identification of molecular signatures based only on the information present in the spatial transcriptomics dataset. We chose herein a reference-free deconvolution algorithm (STdeconvolve) since no JDM single cell/nuclei transcriptomes datasets are publicly available [12] . Reference-free deconvolution prevents us from missing myofibres, and others cell types affected by JDM, that are not present in reference database [27] and enables us to unveil unexpected and novel diversity of JDM associated RNA signatures.

This approach allowed us to identify four physiological and pathological transcriptomic signatures with biological relevance that are heterogeneously distributed within patient’s biopsies. Indeed, in patient presenting with severe and acute NXP2+ JDM, the model of deconvolution identified an almost complete disappearance of the healthy transcriptomic signature in muscle tissue substituted by a prominent pathological myofibrillar stress signature. We observed that this pathological signature within the muscle tissue was preferentially associated with ischaemic changes in myofibres including ischaemic punch-out vacuoles and microinfarcts. *ANKRD1* was the most upregulated gene from the myofibrillar stress signature along with genes encoding for sarcomeric proteins essential for muscle contraction. *ANKRD1* encodes for a pleotropic transcriptional coactivator highly conserved across mammals and is involved in various biological processes, including cardiac muscle development [29] and remodeling following injury or stress [30]. Interactions between this protein and the sarcomeric proteins myopalladin and titin suggest that it is involved in the myofibrillar stretch-sensor system [31]. Its overexpression can lead to heart diseases, such as diastolic dysfunction and heart failure [32]. More recently *ANKRD1* was identified as a mesenchymal-specific transcriptional coactivator driving fibroblast activation in cancer but also in other conditions such as hypertrophic scarring and idiopathic pulmonary fibrosis [33]. *ANKRD1* is also activated in renal ischaemia□reperfusion injury (IRI) models in vivo and in vitro [34]. Altogether, these findings highlight the potential role of *ANKRD1* as a biomarker or therapeutic target for pathological tissue regeneration in JDM. As vasculopathy is central to the pathogenesis of juvenile dermatomyositis [35], the role of *ANKRD1* as a mediator of myofibre response to abnormal blood flow may be further investigated to gain insight into the pathophysiology of myofibre ischaemic changes that occur in severe JDM.

Conversely, in indolent, slowly progressive, JDM, the “muscle remodelling signature” was mostly present. This signature is composed of a mixture of genes encoding for proteins involved in extracellular matrix remodelling, mitochondrial biology, angiogenesis and immune functions. We identified differentially expressed genes that are involved in this pathological signature associated with myofibre changes mostly consisting of a global atrophy. Among these, some genes that have never been described in JDM muscle may be of particular interest. *FSTL1* encodes for a small glycoprotein secreted by skeletal muscle and myocardium and have been implicated in various physiological and pathological processes, including inflammation, wound healing, and cancer progression [36]. The cardioprotective role of FSTL1 has been extensively studied in recent years, although its mechanisms of action remain elusive [37]. In muscle tissue, secretion of FSTL1 by myogenic cells promotes ischaemia-induced revascularisation by activating Akt-endothelial nitric oxide synthase-dependent signaling within endothelial cells [38]. In immune diseases, FSTL1 has a dual function during inflammatory process (anti-inflammatory factor in the acute phase, then pro-inflammatory effect in chronic diseases), probably due to the activation of different signalling pathways [39, 40]. Therefore, FSTL1 may be of interest as a novel non-invasive biomarker or as a new therapeutic target in JDM [41].

The interferon signature was present in the biopsies of all active patients confirming the central role of type I interferon upregulation in JDM pathophysiology. While type I interferon signature has been reported in the muscle of JDM patients by several studies [4, 6, 42], it is worth noting that this signature is very heterogeneously distributed within the biopsies. It suggests that type I interferon may not be homogeneously produced in affected muscle or that type I interferon responding cells may not be homogeneously distributed in the muscle.

A major challenge in the management of JDM patients is the strategy of tapering treatment following good clinical response. Indeed, approximately 50 % of patients will achieve complete clinical remission after first, second- or third-line treatment. Disease relapse can occur even after several years of disease remission [43–45]. Therefore, we need to better understand what the drivers of this risk are and identify biomarkers to better define our tapering strategies in JDM patients. In this study we had the opportunity to study the muscle of a patient performed after 2 years of complete remission following treatment with Ruxolitinib. At the time of her second biopsy, the patient had a normal muscle strength and almost normal muscle biopsy. This was confirmed at the molecular level by the strong reduction of all three pathological signatures compared to the first biopsy. However, only the interferon signature normalised while both “myofibrillar stress” and the muscle remodelling” signatures persisted, suggesting that despite clinical and histological remission, disease signatures persisted.

The persistence of the “muscle remodelling” signature could reflect the process towards complete muscle regeneration to restore a normal muscle function but concomitant “myofibrillar stress” signature suggests that the disease persists and could explain the risk of disease relapse even after a long period of clinical remission [43]. The complete disappearance of the interferon signature is consistent with many previous publications linking interferon protein and signature to disease activity [46].

This study has several limitations. First, we did not combine single-nucleus RNA sequencing with spatial transcriptomics from patient muscle biopsies to map cell-type-specific drivers underlying JDM pathogenesis. Second, our results should be confirmed in a larger cohort of JDM muscle tissues with different MSA subtypes and outcomes to refine correlations between histological/transcriptomic data and to decipher the spatial/temporal chronology of the muscle lesions. Third, this study is basically descriptive, and the observed correlations should not imply causality without complementary investigations.

This study represents the first application of spatial transcriptomics to JDM muscle biopsies using the Visium Spatial Gene Expression technology unveiling the disappearance of normal mitochondrial muscle signature substituted by pathological signatures depending on clinical severity in JDM. Furthermore, abnormal disease signatures persist while patients are on remission. These findings provide promising insights into possible mechanisms of myofibre injuries and disease relapse after remission in JDM.

## Supporting information

Supplemental date

## Patient, public, involvement and engagement

This research has been carried out on biological samples and clinical data collected from children with JDM. In the development of this body of work, it was presented and discussed at the JDM family day as part of the partnership with the Inflammatory Myopathies interest group (https://www.afm-telethon.fr/fr/vivre-avec-la-maladie/l-afm-telethon-m-accompagne/etre-en-contact-avec-d-autres-familles/groupe-myopathies-inflammatoires). The patients, families, researchers and clinical professionals were in attendance at this meeting.

## Acknowledgements

The authors thank all the Children and their families who have contributed to this research. The authors also thank Dr Jules Gilet who initiated the initial muscle spatial transcriptomic experiment.

## Data availability statement

The data that support the findings of this study are openly available in BioStudies at http://doi.org/studies/ S-BSST1845, reference number S-BSST1845.

## Ethics approval

This study was performed in line with the principles of the Declaration of Helsinki. In accordance with current French legislation and hospital research ethics committee (Approval #12-009 at CPP Ile-de-France IX), parents or legal representatives gave their written informed consent for the participation of the children to the study.

## Supplementary-see supplemental materials

**Supplemental Figure S1.**
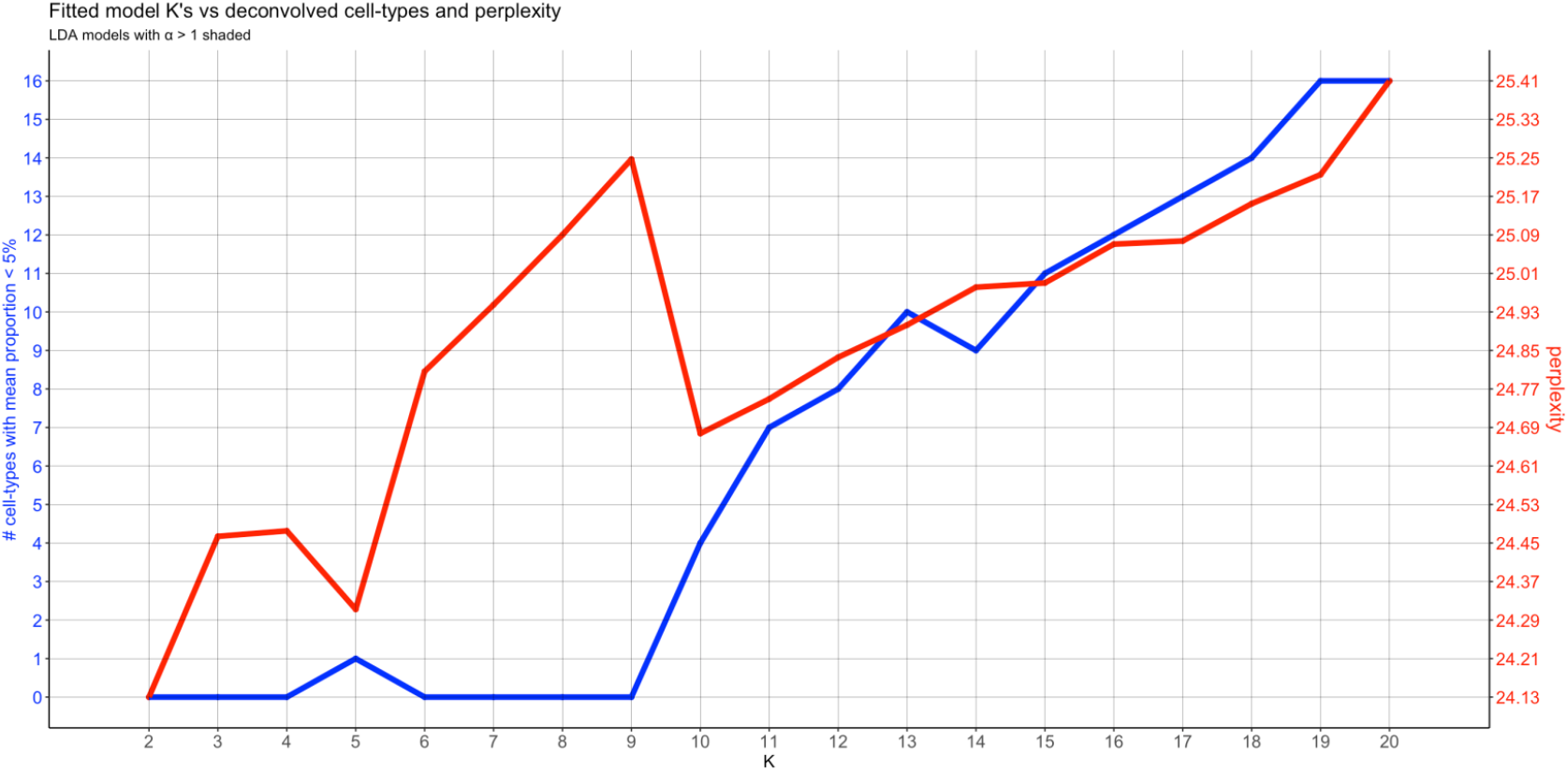

**Supplemental Figure S2.**
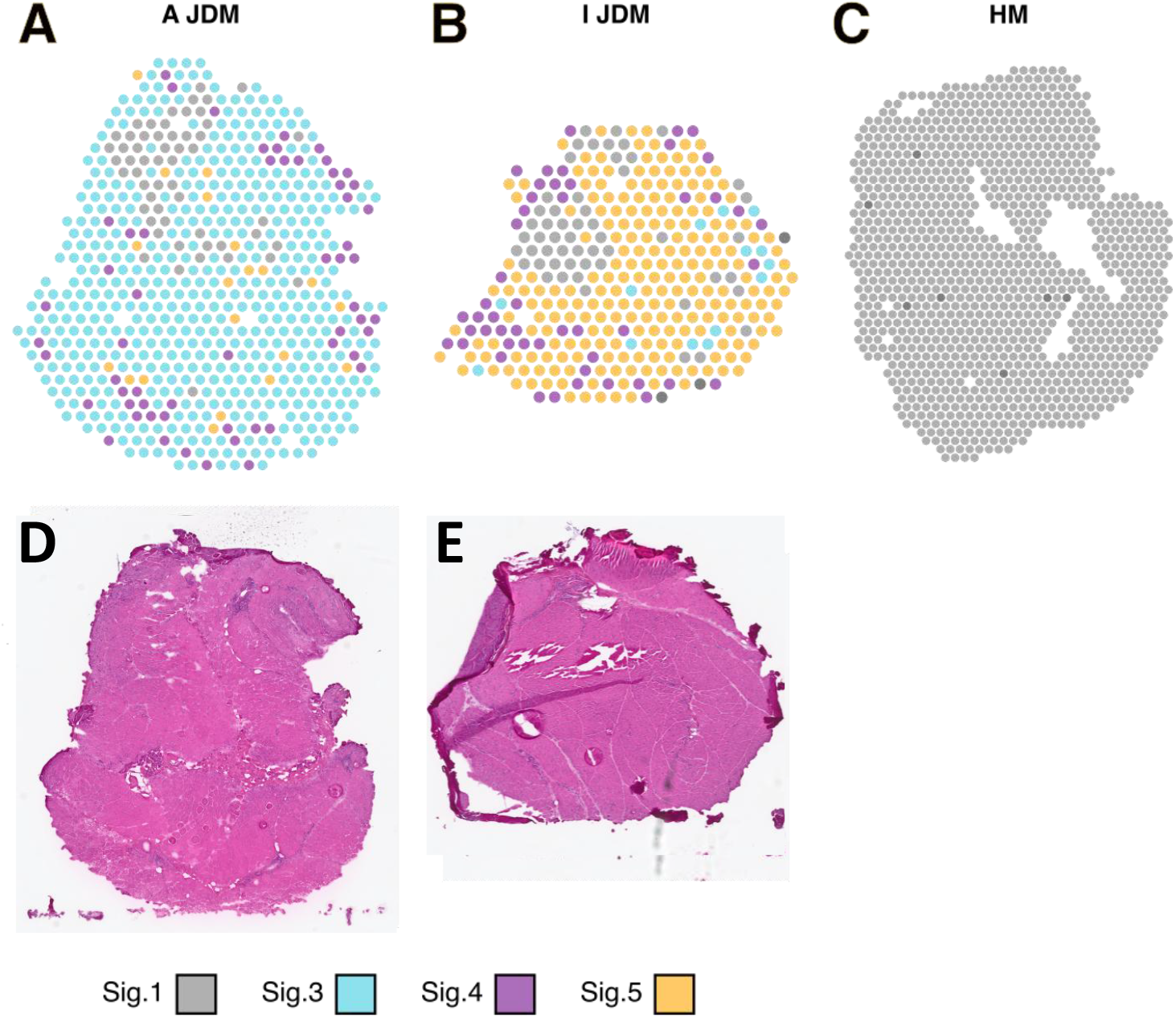

**Supplemental Figure S2.**
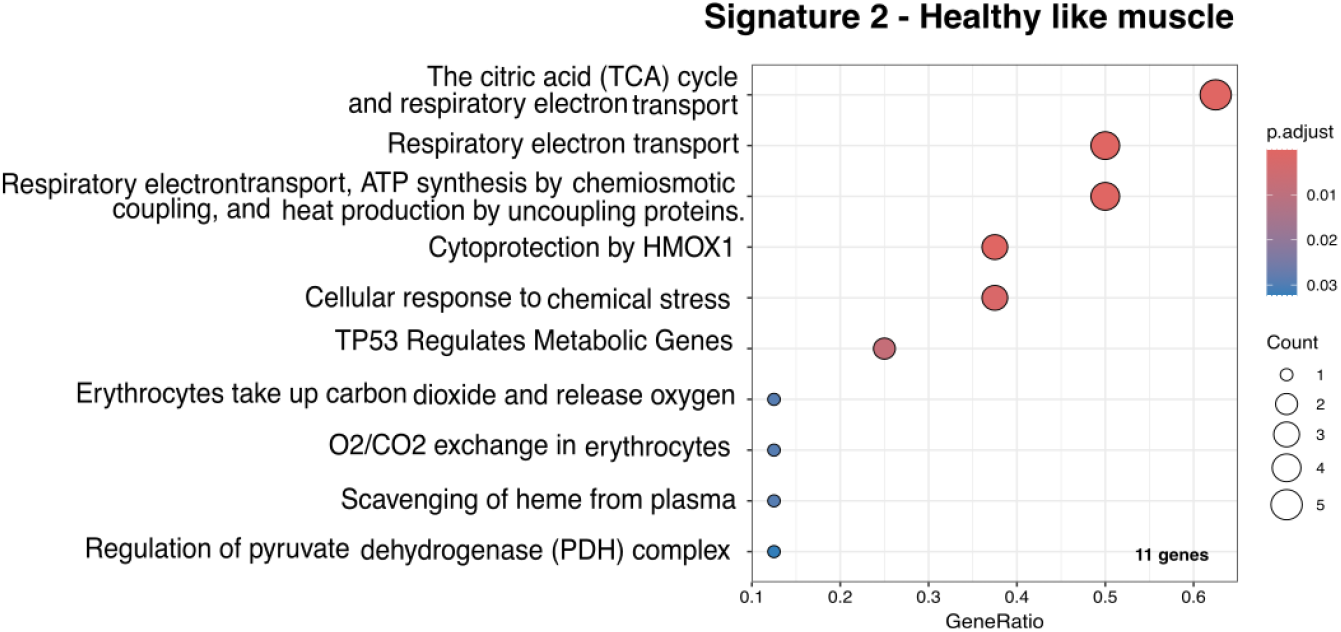

