## Supplemental date for "Muscle spatial transcriptomic reveals heterogeneous profiles in untreated juvenile dermatomyositis and the persistence of pathological signature after remission"

**Tragin et al-Supplementary file**

**Methods**

**Histology, immunohistochemistry, morphometric analyses**

7 μm cryosections of muscle samples (CryoStar NX70, Thermo Fisher Scientific, Waltham, MA, USA) were dried for 30 minutes at room temperature and stored at -80°C before staining. For immunofluorescence staining, sections were thawed for 30 minutes at room temperature and then hydrated with 1X PBS for 10 minutes. They were permeabilized in Triton 0.5% for 5 minutes and then, incubated in 10% BSA (Sigma-Aldrich) for 30 minutes. Sections were incubated with primary antibodies diluted in 1X PBS overnight at 4°C: anti-Laminin rabbit polyclonal antibody (Cat # L9393, 1:500; Sigma-Aldrich) or anti-collagen VI Mouse monoclonal antibody (Cat # MAB1944, 1:1000; Chemicon, Sigma-Aldrich). Then, sections were incubated with appropriate conjugated secondary antibodies for 30 minutes at 37°C and mounted with Fluoromount-G Mounting Medium (Invitrogen). Pictures were acquired using a fluorescent microscope (ZEISS Axio Imager D1). For double immunofluorescence stainings, sections were hydrated with 1X PBS for 10 minutes and then were fixed in 4% PFA for 10 minutes. After this step, the same protocol was applied as above. Sections were incubated with Anti-Laminin rabbit polyclonal antibody (Cat # L9393, 1:500; Sigma-Aldrich) and anti-CD31 endothelial cell Monoclonal Mouse antibody (Cat # M0823, 1:50; Dako, Agilent). Morphometric analysis were performed using a macro developed on Fiji, an open access image processing package based on ImageJ®. This macro allowed to detect and quantify the number of myofibres in sections and to obtain the area or minor diameter values. The other macro used, detected the membrane and vessels associated to each myofibre. To evaluate fibrosis, Fiji was used to detect the whole section and then measure the Collagen VI area fraction.

**Supplemental Table S1** List of gene markers for each of 5 signatures. Reference-free deconvolution provided an expression matrix of genes in each signature. For each signature, marker genes were extracted based on gene expression levels and a fold change was calculated as described in Miller *et al.* (2022).

| Ensemble identifier | Gene marker | log2FC | Molecular Signature |
| --- | --- | --- | --- |
| ENSG00000198899 | ATP6 | 3,20 | 1 |
| ENSG00000228253 | ATP8 | 3,02 | 1 |
| ENSG00000198786 | ND5 | 2,47 | 1 |
| ENSG00000198840 | ND3 | 2,42 | 1 |
| ENSG00000004799 | PDK4 | 2,24 | 1 |
| ENSG00000198804 | COX1 | 2,11 | 1 |
| ENSG00000198938 | COX3 | 1,54 | 1 |
| ENSG00000198727 | CYTB | 1,36 | 1 |
| ENSG00000212907 | ND4L | 1,24 | 1 |
| ENSG00000112306 | RPS12 | -1,46 | 1 |
| ENSG00000184076 | UQCR10 | -1,64 | 1 |
| ENSG00000170290 | SLN | -1,77 | 1 |
| ENSG00000101608 | MYL12A | -2,21 | 1 |
| ENSG00000130595 | TNNT3 | -2,29 | 1 |
| ENSG00000129170 | CSRP3 | -2,30 | 1 |
| ENSG00000106211 | HSPB1 | -2,37 | 1 |
| ENSG00000168530 | MYL1 | -2,38 | 1 |
| ENSG00000101470 | TNNC2 | -2,43 | 1 |
| ENSG00000128591 | FLNC | -2,60 | 1 |
| ENSG00000154553 | PDLIM3 | -2,64 | 1 |
| ENSG00000075624 | ACTB | -2,69 | 1 |
| ENSG00000130598 | TNNI2 | -3,19 | 1 |
| ENSG00000140416 | TPM1 | -3,48 | 1 |
| ENSG00000156508 | EEF1A1 | -3,51 | 1 |
| ENSG00000166710 | B2M | -3,66 | 1 |
| ENSG00000180209 | MYL11 | -3,84 | 1 |
| ENSG00000125414 | MYH2 | -4,02 | 1 |
| ENSG00000244734 | HBB | 27,11 | 2 |
| ENSG00000178878 | APOLD1 | 4,39 | 2 |
| ENSG00000198886 | ND4 | 2,69 | 2 |
| ENSG00000212907 | ND4L | 2,52 | 2 |
| ENSG00000198712 | COX2 | 2,09 | 2 |
| ENSG00000198727 | CYTB | 2,02 | 2 |
| ENSG00000182676 | PPP1R27 | 1,60 | 2 |
| ENSG00000185559 | DLK1 | 1,52 | 2 |
| ENSG00000117479 | SLC19A2 | 1,40 | 2 |
| ENSG00000198938 | COX3 | 1,11 | 2 |
| ENSG00000004799 | PDK4 | 1,09 | 2 |
| ENSG00000142089 | IFITM3 | -1,09 | 2 |
| ENSG00000198899 | ATP6 | -1,39 | 2 |
| ENSG00000142156 | COL6A1 | -1,45 | 2 |
| ENSG00000173641 | HSPB7 | -1,46 | 2 |
| ENSG00000101439 | CST3 | -1,62 | 2 |
| ENSG00000204287 | HLA-DRA | -1,68 | 2 |
| ENSG00000106211 | HSPB1 | -1,93 | 2 |
| ENSG00000101470 | TNNC2 | -1,94 | 2 |
| ENSG00000206503 | HLA-A | -2,27 | 2 |
| ENSG00000011465 | DCN | -2,37 | 2 |
| ENSG00000180209 | MYL11 | -2,47 | 2 |
| ENSG00000168530 | MYL1 | -2,58 | 2 |
| ENSG00000154553 | PDLIM3 | -2,61 | 2 |
| ENSG00000204525 | HLA-C | -2,76 | 2 |
| ENSG00000125414 | MYH2 | -2,84 | 2 |
| ENSG00000130595 | TNNT3 | -2,95 | 2 |
| ENSG00000166710 | B2M | -2,98 | 2 |
| ENSG00000170290 | SLN | -3,58 | 2 |
| ENSG00000148677 | ANKRD1 | 8,01 | 3 |
| ENSG00000159251 | ACTC1 | 5,05 | 3 |
| ENSG00000133020 | MYH8 | 4,87 | 3 |
| ENSG00000109063 | MYH3 | 4,14 | 3 |
| ENSG00000138435 | CHRNA1 | 3,91 | 3 |
| ENSG00000165887 | ANKRD2 | 2,95 | 3 |
| ENSG00000101608 | MYL12A | 2,83 | 3 |
| ENSG00000150779 | TIMM8B | 2,63 | 3 |
| ENSG00000170290 | SLN | 2,56 | 3 |
| ENSG00000180209 | MYL11 | 2,53 | 3 |
| ENSG00000086967 | MYBPC2 | 2,49 | 3 |
| ENSG00000106992 | AK1 | 2,43 | 3 |
| ENSG00000168530 | MYL1 | 2,31 | 3 |
| ENSG00000272573 | MUSTN1 | 2,21 | 3 |
| ENSG00000166025 | AMOTL1 | 2,10 | 3 |
| ENSG00000118729 | CASQ2 | 1,91 | 3 |
| ENSG00000140416 | TPM1 | 1,84 | 3 |
| ENSG00000106211 | HSPB1 | 1,82 | 3 |
| ENSG00000133110 | POSTN | 1,71 | 3 |
| ENSG00000267855 | NDUFA7 | 1,68 | 3 |
| ENSG00000126709 | IFI6 | 1,65 | 3 |
| ENSG00000109061 | MYH1 | 1,62 | 3 |
| ENSG00000154553 | PDLIM3 | 1,35 | 3 |
| ENSG00000101470 | TNNC2 | 1,26 | 3 |
| ENSG00000128591 | FLNC | 1,16 | 3 |
| ENSG00000130595 | TNNT3 | 1,05 | 3 |
| ENSG00000125414 | MYH2 | 1,02 | 3 |
| ENSG00000101439 | CST3 | -1,07 | 3 |
| ENSG00000163430 | FSTL1 | -1,18 | 3 |
| ENSG00000142156 | COL6A1 | -1,27 | 3 |
| ENSG00000008517 | IL32 | -1,31 | 3 |
| ENSG00000111341 | MGP | -1,59 | 3 |
| ENSG00000198899 | ATP6 | -1,59 | 3 |
| ENSG00000198786 | ND5 | -1,60 | 3 |
| ENSG00000198712 | COX2 | -1,74 | 3 |
| ENSG00000198804 | COX1 | -1,79 | 3 |
| ENSG00000026025 | VIM | -1,82 | 3 |
| ENSG00000163453 | IGFBP7 | -1,82 | 3 |
| ENSG00000198840 | ND3 | -2,14 | 3 |
| ENSG00000011465 | DCN | -2,26 | 3 |
| ENSG00000148180 | GSN | -2,26 | 3 |
| ENSG00000142089 | IFITM3 | -2,49 | 3 |
| ENSG00000198938 | COX3 | -3,13 | 3 |
| ENSG00000198727 | CYTB | -3,68 | 3 |
| ENSG00000198886 | ND4 | -5,33 | 3 |
| ENSG00000211592 | IGKC | 4,90 | 4 |
| ENSG00000173369 | C1QB | 4,71 | 4 |
| ENSG00000107796 | ACTA2 | 4,10 | 4 |
| ENSG00000187608 | ISG15 | 4,06 | 4 |
| ENSG00000100600 | LGMN | 3,79 | 4 |
| ENSG00000185885 | IFITM1 | 3,61 | 4 |
| ENSG00000115415 | STAT1 | 3,57 | 4 |
| ENSG00000119917 | IFIT3 | 3,46 | 4 |
| ENSG00000182326 | C1S | 3,42 | 4 |
| ENSG00000111341 | MGP | 3,40 | 4 |
| ENSG00000185745 | IFIT1 | 3,36 | 4 |
| ENSG00000019582 | CD74 | 3,18 | 4 |
| ENSG00000204287 | HLA-DRA | 3,15 | 4 |
| ENSG00000165949 | IFI27 | 3,01 | 4 |
| ENSG00000108821 | COL1A1 | 3,00 | 4 |
| ENSG00000163191 | S100A11 | 2,96 | 4 |
| ENSG00000087245 | MMP2 | 2,88 | 4 |
| ENSG00000075624 | ACTB | 2,76 | 4 |
| ENSG00000196262 | PPIA | 2,60 | 4 |
| ENSG00000231389 | HLA-DPA1 | 2,60 | 4 |
| ENSG00000165516 | KLHDC2 | 2,59 | 4 |
| ENSG00000142089 | IFITM3 | 2,53 | 4 |
| ENSG00000175899 | A2M | 2,52 | 4 |
| ENSG00000149131 | SERPING1 | 2,48 | 4 |
| ENSG00000206503 | HLA-A | 2,46 | 4 |
| ENSG00000137965 | IFI44 | 2,41 | 4 |
| ENSG00000026025 | VIM | 2,38 | 4 |
| ENSG00000108679 | LGALS3BP | 2,38 | 4 |
| ENSG00000130303 | BST2 | 2,37 | 4 |
| ENSG00000126709 | IFI6 | 2,35 | 4 |
| ENSG00000234745 | HLA-B | 2,32 | 4 |
| ENSG00000133110 | POSTN | 2,29 | 4 |
| ENSG00000129538 | RNASE1 | 2,29 | 4 |
| ENSG00000113140 | SPARC | 2,28 | 4 |
| ENSG00000166710 | B2M | 2,27 | 4 |
| ENSG00000142156 | COL6A1 | 2,26 | 4 |
| ENSG00000135404 | CD63 | 2,21 | 4 |
| ENSG00000164692 | COL1A2 | 2,16 | 4 |
| ENSG00000204525 | HLA-C | 2,16 | 4 |
| ENSG00000163453 | IGFBP7 | 2,16 | 4 |
| ENSG00000034510 | TMSB10 | 2,16 | 4 |
| ENSG00000184009 | ACTG1 | 2,11 | 4 |
| ENSG00000159403 | C1R | 1,99 | 4 |
| ENSG00000123416 | TUBA1B | 1,97 | 4 |
| ENSG00000112695 | COX7A2 | 1,87 | 4 |
| ENSG00000204291 | COL15A1 | 1,86 | 4 |
| ENSG00000160932 | LY6E | 1,86 | 4 |
| ENSG00000135046 | ANXA1 | 1,85 | 4 |
| ENSG00000187498 | COL4A1 | 1,78 | 4 |
| ENSG00000185088 | RPS27L | 1,78 | 4 |
| ENSG00000163430 | FSTL1 | 1,75 | 4 |
| ENSG00000107562 | CXCL12 | 1,72 | 4 |
| ENSG00000115414 | FN1 | 1,70 | 4 |
| ENSG00000138615 | CILP | 1,68 | 4 |
| ENSG00000197747 | S100A10 | 1,62 | 4 |
| ENSG00000156508 | EEF1A1 | 1,59 | 4 |
| ENSG00000163513 | TGFBR2 | 1,56 | 4 |
| ENSG00000159377 | PSMB4 | 1,51 | 4 |
| ENSG00000134871 | COL4A2 | 1,50 | 4 |
| ENSG00000168542 | COL3A1 | 1,49 | 4 |
| ENSG00000101439 | CST3 | 1,43 | 4 |
| ENSG00000197956 | S100A6 | 1,41 | 4 |
| ENSG00000163520 | FBLN2 | 1,40 | 4 |
| ENSG00000112306 | RPS12 | 1,39 | 4 |
| ENSG00000185559 | DLK1 | 1,38 | 4 |
| ENSG00000166147 | FBN1 | 1,30 | 4 |
| ENSG00000086967 | MYBPC2 | 1,30 | 4 |
| ENSG00000115541 | HSPE1 | 1,11 | 4 |
| ENSG00000166025 | AMOTL1 | 1,02 | 4 |
| ENSG00000159251 | ACTC1 | -1,05 | 4 |
| ENSG00000140416 | TPM1 | -1,08 | 4 |
| ENSG00000154553 | PDLIM3 | -1,15 | 4 |
| ENSG00000106211 | HSPB1 | -1,15 | 4 |
| ENSG00000129170 | CSRP3 | -1,26 | 4 |
| ENSG00000165887 | ANKRD2 | -1,55 | 4 |
| ENSG00000101608 | MYL12A | -1,65 | 4 |
| ENSG00000004799 | PDK4 | -1,73 | 4 |
| ENSG00000130595 | TNNT3 | -1,81 | 4 |
| ENSG00000198712 | COX2 | -2,07 | 4 |
| ENSG00000130598 | TNNI2 | -2,13 | 4 |
| ENSG00000198899 | ATP6 | -2,21 | 4 |
| ENSG00000125414 | MYH2 | -2,23 | 4 |
| ENSG00000198786 | ND5 | -2,32 | 4 |
| ENSG00000180209 | MYL11 | -2,38 | 4 |
| ENSG00000101470 | TNNC2 | -2,47 | 4 |
| ENSG00000198727 | CYTB | -3,10 | 4 |
| ENSG00000168530 | MYL1 | -3,17 | 4 |
| ENSG00000198804 | COX1 | -3,21 | 4 |
| ENSG00000198886 | ND4 | -3,53 | 4 |
| ENSG00000198840 | ND3 | -4,76 | 4 |
| ENSG00000198938 | COX3 | -5,09 | 4 |
| ENSG00000197766 | CFD | 5,66 | 5 |
| ENSG00000008517 | IL32 | 3,09 | 5 |
| ENSG00000148180 | GSN | 2,96 | 5 |
| ENSG00000214548 | MEG3 | 2,88 | 5 |
| ENSG00000158710 | TAGLN2 | 2,70 | 5 |
| ENSG00000011465 | DCN | 2,65 | 5 |
| ENSG00000129170 | CSRP3 | 2,56 | 5 |
| ENSG00000130598 | TNNI2 | 2,49 | 5 |
| ENSG00000125414 | MYH2 | 2,38 | 5 |
| ENSG00000136521 | NDUFB5 | 2,19 | 5 |
| ENSG00000173641 | HSPB7 | 2,16 | 5 |
| ENSG00000147684 | NDUFB9 | 2,11 | 5 |
| ENSG00000130595 | TNNT3 | 2,07 | 5 |
| ENSG00000163520 | FBLN2 | 2,03 | 5 |
| ENSG00000101470 | TNNC2 | 1,87 | 5 |
| ENSG00000135046 | ANXA1 | 1,84 | 5 |
| ENSG00000163359 | COL6A3 | 1,80 | 5 |
| ENSG00000159403 | C1R | 1,74 | 5 |
| ENSG00000113296 | THBS4 | 1,74 | 5 |
| ENSG00000204291 | COL15A1 | 1,74 | 5 |
| ENSG00000108561 | C1QBP | 1,69 | 5 |
| ENSG00000163430 | FSTL1 | 1,67 | 5 |
| ENSG00000117479 | SLC19A2 | 1,64 | 5 |
| ENSG00000106588 | PSMA2 | 1,61 | 5 |
| ENSG00000154553 | PDLIM3 | 1,60 | 5 |
| ENSG00000134871 | COL4A2 | 1,58 | 5 |
| ENSG00000168542 | COL3A1 | 1,57 | 5 |
| ENSG00000166147 | FBN1 | 1,52 | 5 |
| ENSG00000138615 | CILP | 1,47 | 5 |
| ENSG00000149131 | SERPING1 | 1,44 | 5 |
| ENSG00000115541 | HSPE1 | 1,39 | 5 |
| ENSG00000101439 | CST3 | 1,33 | 5 |
| ENSG00000178209 | PLEC | 1,32 | 5 |
| ENSG00000128591 | FLNC | 1,32 | 5 |
| ENSG00000163513 | TGFBR2 | 1,25 | 5 |
| ENSG00000267855 | NDUFA7 | 1,23 | 5 |
| ENSG00000197956 | S100A6 | 1,20 | 5 |
| ENSG00000184076 | UQCR10 | 1,15 | 5 |
| ENSG00000026025 | VIM | 1,05 | 5 |
| ENSG00000155096 | AZIN1 | 1,02 | 5 |
| ENSG00000198786 | ND5 | -1,09 | 5 |
| ENSG00000234745 | HLA-B | -1,17 | 5 |
| ENSG00000198899 | ATP6 | -1,38 | 5 |
| ENSG00000198804 | COX1 | -1,54 | 5 |
| ENSG00000206503 | HLA-A | -1,84 | 5 |
| ENSG00000198727 | CYTB | -2,01 | 5 |
| ENSG00000198712 | COX2 | -2,21 | 5 |
| ENSG00000198886 | ND4 | -2,73 | 5 |

**Supplemental Table S2: Indolent and acute JDM muscles display distinct transcriptomic signatures (all results).** All significantly enriched Reactome pathways in each signature. Figure 3 provides only the 10 first pathways.

| Molecular signature | ID | Description | GeneRatio | BgRatio | pvalue | p.adjust | qvalue | geneID | Count |
| --- | --- | --- | --- | --- | --- | --- | --- | --- | --- |
| 1 | R-HSA-1428517 | The citric acid (TCA) cycle and respiratory electron transport | 8/8 | 178/11009 | 3,99E-15 | 6,79E-14 | 4,20E-15 | ATP6/ATP8/ND5/ND3/PDK4/COX1/COX3/CYTB | 8 |
| 1 | R-HSA-163200 | Respiratory electron transport, ATP synthesis by chemiosmotic coupling, and heat production by uncoupling proteins. | 7/8 | 127/11009 | 1,82E-13 | 1,55E-12 | 9,60E-14 | ATP6/ATP8/ND5/ND3/COX1/COX3/CYTB | 7 |
| 1 | R-HSA-611105 | Respiratory electron transport | 5/8 | 103/11009 | 3,56E-09 | 2,02E-08 | 1,25E-09 | ND5/ND3/COX1/COX3/CYTB | 5 |
| 1 | R-HSA-163210 | Formation of ATP by chemiosmotic coupling | 2/8 | 18/11009 | 7,03E-05 | 2,99E-04 | 1,85E-05 | ATP6/ATP8 | 2 |
| 1 | R-HSA-8949613 | Cristae formation | 2/8 | 31/11009 | 2,13E-04 | 7,23E-04 | 4,48E-05 | ATP6/ATP8 | 2 |
| 1 | R-HSA-6799198 | Complex I biogenesis | 2/8 | 57/11009 | 7,23E-04 | 2,05E-03 | 1,27E-04 | ND5/ND3 | 2 |
| 1 | R-HSA-9707564 | Cytoprotection by HMOX1 | 2/8 | 64/11009 | 9,11E-04 | 2,21E-03 | 1,37E-04 | COX1/COX3 | 2 |
| 1 | R-HSA-5628897 | TP53 Regulates Metabolic Genes | 2/8 | 87/11009 | 1,68E-03 | 3,56E-03 | 2,21E-04 | COX1/COX3 | 2 |
| 1 | R-HSA-1592230 | Mitochondrial biogenesis | 2/8 | 95/11009 | 1,99E-03 | 3,77E-03 | 2,33E-04 | ATP6/ATP8 | 2 |
| 1 | R-HSA-9711123 | Cellular response to chemical stress | 2/8 | 215/11009 | 9,84E-03 | 1,67E-02 | 1,04E-03 | COX1/COX3 | 2 |
| 1 | R-HSA-204174 | Regulation of pyruvate dehydrogenase (PDH) complex | 1/8 | 16/11009 | 1,16E-02 | 1,79E-02 | 1,11E-03 | PDK4 | 1 |
| 1 | R-HSA-1852241 | Organelle biogenesis and maintenance | 2/8 | 296/11009 | 1,81E-02 | 2,57E-02 | 1,59E-03 | ATP6/ATP8 | 2 |
| 1 | R-HSA-70268 | Pyruvate metabolism | 1/8 | 31/11009 | 2,23E-02 | 2,92E-02 | 1,81E-03 | PDK4 | 1 |
| 1 | R-HSA-3700989 | Transcriptional Regulation by TP53 | 2/8 | 362/11009 | 2,65E-02 | 3,22E-02 | 1,99E-03 | COX1/COX3 | 2 |
| 1 | R-HSA-5362517 | Signaling by Retinoic Acid | 1/8 | 43/11009 | 3,08E-02 | 3,49E-02 | 2,16E-03 | PDK4 | 1 |
| 1 | R-HSA-71406 | Pyruvate metabolism and Citric Acid (TCA) cycle | 1/8 | 55/11009 | 3,93E-02 | 4,17E-02 | 2,58E-03 | PDK4 | 1 |
| 2 | R-HSA-1428517 | The citric acid (TCA) cycle and respiratory electron transport | 5/8 | 178/11009 | 5,63E-08 | 1,58E-06 | 4,74E-07 | ND4/COX2/CYTB/COX3/PDK4 | 5 |
| 2 | R-HSA-611105 | Respiratory electron transport | 4/8 | 103/11009 | 4,92E-07 | 6,88E-06 | 2,07E-06 | ND4/COX2/CYTB/COX3 | 4 |
| 2 | R-HSA-163200 | Respiratory electron transport, ATP synthesis by chemiosmotic coupling, and heat production by uncoupling proteins. | 4/8 | 127/11009 | 1,14E-06 | 1,06E-05 | 3,20E-06 | ND4/COX2/CYTB/COX3 | 4 |
| 2 | R-HSA-9707564 | Cytoprotection by HMOX1 | 3/8 | 64/11009 | 1,03E-05 | 7,20E-05 | 2,16E-05 | HBB/COX2/COX3 | 3 |
| 2 | R-HSA-9711123 | Cellular response to chemical stress | 3/8 | 215/11009 | 3,83E-04 | 2,14E-03 | 6,44E-04 | HBB/COX2/COX3 | 3 |
| 2 | R-HSA-5628897 | TP53 Regulates Metabolic Genes | 2/8 | 87/11009 | 1,68E-03 | 7,82E-03 | 2,35E-03 | COX2/COX3 | 2 |
| 2 | R-HSA-1237044 | Erythrocytes take up carbon dioxide and release oxygen | 1/8 | 13/11009 | 9,41E-03 | 2,93E-02 | 8,81E-03 | HBB | 1 |
| 2 | R-HSA-1480926 | O2/CO2 exchange in erythrocytes | 1/8 | 13/11009 | 9,41E-03 | 2,93E-02 | 8,81E-03 | HBB | 1 |
| 2 | R-HSA-2168880 | Scavenging of heme from plasma | 1/8 | 13/11009 | 9,41E-03 | 2,93E-02 | 8,81E-03 | HBB | 1 |
| 2 | R-HSA-204174 | Regulation of pyruvate dehydrogenase (PDH) complex | 1/8 | 16/11009 | 1,16E-02 | 3,24E-02 | 9,74E-03 | PDK4 | 1 |
| 2 | R-HSA-9613829 | Chaperone Mediated Autophagy | 1/8 | 22/11009 | 1,59E-02 | 4,04E-02 | 1,22E-02 | HBB | 1 |
| 2 | R-HSA-2122948 | Activated NOTCH1 Transmits Signal to the Nucleus | 1/8 | 31/11009 | 2,23E-02 | 4,81E-02 | 1,45E-02 | DLK1 | 1 |
| 2 | R-HSA-70268 | Pyruvate metabolism | 1/8 | 31/11009 | 2,23E-02 | 4,81E-02 | 1,45E-02 | PDK4 | 1 |
| 2 | R-HSA-9615710 | Late endosomal microautophagy | 1/8 | 35/11009 | 2,52E-02 | 4,94E-02 | 1,49E-02 | HBB | 1 |
| 2 | R-HSA-3700989 | Transcriptional Regulation by TP53 | 2/8 | 362/11009 | 2,65E-02 | 4,94E-02 | 1,49E-02 | COX2/COX3 | 2 |
| 3 | R-HSA-397014 | Muscle contraction | 12/22 | 205/11009 | 6,92E-16 | 4,22E-14 | 3,20E-14 | ACTC1/MYH8/MYH3/MYL12A/SLN/MYL11/MYBPC2/MYL1/CASQ2/TPM1/TNNC2/TNNT3 | 12 |
| 3 | R-HSA-390522 | Striated Muscle Contraction | 8/22 | 36/11009 | 1,76E-15 | 5,36E-14 | 4,07E-14 | ACTC1/MYH8/MYH3/MYBPC2/MYL1/TPM1/TNNC2/TNNT3 | 8 |
| 3 | R-HSA-445355 | Smooth Muscle Contraction | 3/22 | 45/11009 | 9,31E-05 | 1,89E-03 | 1,44E-03 | MYL12A/MYL11/TPM1 | 3 |
| 4 | R-HSA-909733 | Interferon alpha/beta signaling | 12/61 | 77/11009 | 7,44E-15 | 3,27E-12 | 2,41E-12 | ISG15/IFITM1/STAT1/IFIT3/IFIT1/IFI27/IFITM3/HLA-A /BST2/ IFI6 / HLA-B/HLA-C | 12 |
| 4 | R-HSA-913531 | Interferon Signaling | 16/61 | 269/11009 | 7,84E-13 | 1,72E-10 | 1,27E-10 | ISG15/IFITM1/STAT1/IFIT3/IFIT1/HLA-DRA/IFI27/HLA-DPA1/IFITM3/ HLA-A/BST2/IFI6/HLA-B/B2M/HLA-C/TUBA1B | 16 |
| 4 | R-HSA-1474228 | Degradation of the extracellular matrix | 11/61 | 140/11009 | 2,30E-10 | 3,37E-08 | 2,48E-08 | COL1A1/MMP2/A2M/COL6A1/COL1A2/COL15A1/COL4A1/FN1/COL4A2/COL3A1/FBN1 | 11 |
| 4 | R-HSA-1442490 | Collagen degradation | 8/61 | 64/11009 | 1,92E-09 | 2,11E-07 | 1,55E-07 | COL1A1/MMP2/COL6A1/COL1A2/COL15A1/COL4A1/COL4A2/COL3A1 | 8 |
| 4 | R-HSA-8948216 | Collagen chain trimerization | 7/61 | 44/11009 | 3,67E-09 | 3,23E-07 | 2,37E-07 | COL1A1/COL6A1/COL1A2/COL15A1/COL4A1/COL4A2/COL3A1 | 7 |
| 4 | R-HSA-1474244 | Extracellular matrix organization | 13/61 | 300/11009 | 7,16E-09 | 4,88E-07 | 3,59E-07 | COL1A1/MMP2/A2M/SPARC/COL6A1/COL1A2/COL15A1/COL4A1/FN1/COL4A2/COL3A1/FBLN2/FBN1 | 13 |
| 4 | R-HSA-3000178 | ECM proteoglycans | 8/61 | 76/11009 | 7,77E-09 | 4,88E-07 | 3,59E-07 | COL1A1/SPARC/COL6A1/COL1A2/COL4A1/FN1/COL4A2/COL3A1 | 8 |
| 4 | R-HSA-216083 | Integrin cell surface interactions | 8/61 | 85/11009 | 1,91E-08 | 1,05E-06 | 7,70E-07 | COL1A1/COL6A1/COL1A2/COL4A1/FN1/COL4A2/COL3A1/FBN1 | 8 |
| 4 | R-HSA-2022090 | Assembly of collagen fibrils and other multimeric structures | 7/61 | 61/11009 | 3,89E-08 | 1,90E-06 | 1,40E-06 | COL1A1/COL6A1/COL1A2/COL15A1/COL4A1/COL4A2/COL3A1 | 7 |
| 4 | R-HSA-3000480 | Scavenging by Class A Receptors | 5/61 | 19/11009 | 4,84E-08 | 2,13E-06 | 1,56E-06 | COL1A1/COL1A2/COL4A1/COL4A2/COL3A1 | 5 |
| 4 | R-HSA-1650814 | Collagen biosynthesis and modifying enzymes | 7/61 | 67/11009 | 7,55E-08 | 3,02E-06 | 2,22E-06 | COL1A1/COL6A1/COL1A2/COL15A1/COL4A1/COL4A2/COL3A1 | 7 |
| 4 | R-HSA-2173782 | Binding and Uptake of Ligands by Scavenger Receptors | 6/61 | 42/11009 | 1,01E-07 | 3,71E-06 | 2,72E-06 | COL1A1/SPARC/COL1A2/COL4A1/COL4A2/COL3A1 | 6 |
| 4 | R-HSA-1236977 | Endosomal/Vacuolar pathway | 4/61 | 11/11009 | 2,73E-07 | 9,26E-06 | 6,80E-06 | HLA-A/HLA-B/B2M/HLA-C | 4 |
| 4 | R-HSA-1474290 | Collagen formation | 7/61 | 90/11009 | 5,87E-07 | 1,85E-05 | 1,36E-05 | COL1A1/COL6A1/COL1A2/COL15A1/COL4A1/COL4A2/COL3A1 | 7 |
| 4 | R-HSA-198933 | Immunoregulatory interactions between a Lymphoid and a non-Lymphoid cell | 8/61 | 133/11009 | 6,32E-07 | 1,85E-05 | 1,36E-05 | IFITM1/COL1A1/HLA-A/HLA-B/B2M/COL1A2/HLA-C/COL3A1 | 8 |
| 4 | R-HSA-3000171 | Non-integrin membrane-ECM interactions | 6/61 | 59/11009 | 8,07E-07 | 2,22E-05 | 1,63E-05 | COL1A1/COL1A2/COL4A1/FN1/COL4A2/COL3A1 | 6 |
| 4 | R-HSA-877300 | Interferon gamma signaling | 7/61 | 96/11009 | 9,13E-07 | 2,36E-05 | 1,74E-05 | STAT1/HLA-DRA/HLA-DPA1/HLA-A/ HLA-B/B2M/HLA-C | 7 |
| 4 | R-HSA-2214320 | Anchoring fibril formation | 4/61 | 15/11009 | 1,11E-06 | 2,72E-05 | 2,00E-05 | COL1A1/COL1A2/COL4A1/COL4A2 | 4 |
| 4 | R-HSA-2243919 | Crosslinking of collagen fibrils | 4/61 | 18/11009 | 2,46E-06 | 5,70E-05 | 4,19E-05 | COL1A1/COL1A2/COL4A1/COL4A2 | 4 |
| 4 | R-HSA-114608 | Platelet degranulation | 7/61 | 129/11009 | 6,64E-06 | 1,46E-04 | 1,07E-04 | PPIA/A2M/SERPING1/LGALS3BP/SPARC/CD63/FN1 | 7 |
| 4 | R-HSA-76005 | Response to elevated platelet cytosolic Ca2+ | 7/61 | 134/11009 | 8,54E-06 | 1,79E-04 | 1,31E-04 | PPIA/A2M/SERPING1/LGALS3BP/SPARC/CD63/FN1 | 7 |
| 4 | R-HSA-76002 | Platelet activation, signaling and aggregation | 9/61 | 263/11009 | 1,29E-05 | 2,59E-04 | 1,90E-04 | COL1A1/PPIA/A2M/SERPING1/LGALS3BP/SPARC/CD63/COL1A2/FN1 | 9 |
| 4 | R-HSA-3000170 | Syndecan interactions | 4/61 | 27/11009 | 1,36E-05 | 2,60E-04 | 1,91E-04 | COL1A1/COL1A2/FN1/COL3A1 | 4 |
| 4 | R-HSA-9679506 | SARS-CoV Infections | 11/61 | 414/11009 | 1,43E-05 | 2,61E-04 | 1,92E-04 | ISG15/STAT1/PPIA/HLA-A/BST2/HLA-B/B2M/HLA-C/RPS27L/EEF1A1/RPS12 | 11 |
| 4 | R-HSA-9705683 | SARS-CoV-2-host interactions | 8/61 | 203/11009 | 1,48E-05 | 2,61E-04 | 1,92E-04 | ISG15/STAT1/HLA-A/HLA-B/B2M/HLA-C/RPS27L/RPS12 | 8 |
| 4 | R-HSA-186797 | Signaling by PDGF | 5/61 | 58/11009 | 1,62E-05 | 2,74E-04 | 2,01E-04 | STAT1/COL6A1/COL4A1/COL4A2/COL3A1 | 5 |
| 4 | R-HSA-983170 | Antigen Presentation: Folding, assembly and peptide loading of class I MHC | 4/61 | 29/11009 | 1,83E-05 | 2,98E-04 | 2,19E-04 | HLA-A/HLA-B/B2M/HLA-C | 4 |
| 4 | R-HSA-8874081 | MET activates PTK2 signaling | 4/61 | 30/11009 | 2,10E-05 | 3,30E-04 | 2,42E-04 | COL1A1/COL1A2/FN1/COL3A1 | 4 |
| 4 | R-HSA-6785807 | Interleukin-4 and Interleukin-13 signaling | 6/61 | 108/11009 | 2,78E-05 | 4,21E-04 | 3,10E-04 | STAT1/MMP2/VIM/COL1A2/ANXA1/FN1 | 6 |
| 4 | R-HSA-166786 | Creation of C4 and C2 activators | 3/61 | 14/11009 | 5,64E-05 | 8,27E-04 | 6,08E-04 | C1QB/C1S/C1R | 3 |
| 4 | R-HSA-381426 | Regulation of Insulin-like Growth Factor (IGF) transport and uptake by Insulin-like Growth Factor Binding Proteins (IGFBPs) | 6/61 | 125/11009 | 6,33E-05 | 8,99E-04 | 6,60E-04 | MMP2/IGFBP7/FSTL1/FN1/CST3/FBN1 | 6 |
| 4 | R-HSA-9705671 | SARS-CoV-2 activates/modulates innate and adaptive immune responses | 6/61 | 126/11009 | 6,62E-05 | 9,10E-04 | 6,68E-04 | ISG15/STAT1/HLA-A/HLA-B/B2M/HLA-C | 6 |
| 4 | R-HSA-8875878 | MET promotes cell motility | 4/61 | 41/11009 | 7,41E-05 | 9,88E-04 | 7,26E-04 | COL1A1/COL1A2/FN1/COL3A1 | 4 |
| 4 | R-HSA-419037 | NCAM1 interactions | 4/61 | 42/11009 | 8,16E-05 | 1,06E-03 | 7,75E-04 | COL6A1/COL4A1/COL4A2/COL3A1 | 4 |
| 4 | R-HSA-977606 | Regulation of Complement cascade | 4/61 | 47/11009 | 1,27E-04 | 1,60E-03 | 1,18E-03 | C1QB/C1S/SERPING1/C1R | 4 |
| 4 | R-HSA-1236974 | ER-Phagosome pathway | 5/61 | 90/11009 | 1,35E-04 | 1,66E-03 | 1,22E-03 | HLA-A/HLA-B/B2M/HLA-C/PSMB4 | 5 |
| 4 | R-HSA-9692914 | SARS-CoV-1-host interactions | 5/61 | 96/11009 | 1,84E-04 | 2,18E-03 | 1,60E-03 | PPIA/BST2/RPS27L/EEF1A1/RPS12 | 5 |
| 4 | R-HSA-9694516 | SARS-CoV-2 Infection | 8/61 | 299/11009 | 2,27E-04 | 2,61E-03 | 1,92E-03 | ISG15/STAT1/HLA-A/HLA-B/B2M/HLA-C/RPS27L/RPS12 | 8 |
| 4 | R-HSA-9613829 | Chaperone Mediated Autophagy | 3/61 | 22/11009 | 2,31E-04 | 2,61E-03 | 1,92E-03 | VIM/RNASE1/EEF1A1 | 3 |
| 4 | R-HSA-166663 | Initial triggering of complement | 3/61 | 23/11009 | 2,65E-04 | 2,91E-03 | 2,14E-03 | C1QB/C1S/C1R | 3 |
| 4 | R-HSA-1236975 | Antigen processing-Cross presentation | 5/61 | 105/11009 | 2,79E-04 | 3,00E-03 | 2,20E-03 | HLA-A/HLA-B/B2M/HLA-C/PSMB4 | 5 |
| 4 | R-HSA-166658 | Complement cascade | 4/61 | 58/11009 | 2,89E-04 | 3,03E-03 | 2,23E-03 | C1QB/C1S/SERPING1/C1R | 4 |
| 4 | R-HSA-8957275 | Post-translational protein phosphorylation | 5/61 | 108/11009 | 3,18E-04 | 3,26E-03 | 2,39E-03 | IGFBP7/FSTL1/FN1/CST3/FBN1 | 5 |
| 4 | R-HSA-375165 | NCAM signaling for neurite out-growth | 4/61 | 63/11009 | 3,98E-04 | 3,98E-03 | 2,92E-03 | COL6A1/COL4A1/COL4A2/COL3A1 | 4 |
| 4 | R-HSA-2132295 | MHC class II antigen presentation | 5/61 | 123/11009 | 5,79E-04 | 5,66E-03 | 4,16E-03 | LGMN/CD74/HLA-DRA/HLA-DPA1/TUBA1B | 5 |
| 4 | R-HSA-9006936 | Signaling by TGFB family members | 5/61 | 124/11009 | 6,01E-04 | 5,75E-03 | 4,22E-03 | STAT1/COL1A2/FSTL1/TGFBR2/FBN1 | 5 |
| 4 | R-HSA-5626467 | RHO GTPases activate IQGAPs | 3/61 | 32/11009 | 7,16E-04 | 6,70E-03 | 4,92E-03 | ACTB/ACTG1/TUBA1B | 3 |
| 4 | R-HSA-6806834 | Signaling by MET | 4/61 | 79/11009 | 9,40E-04 | 8,47E-03 | 6,22E-03 | COL1A1/COL1A2/FN1/COL3A1 | 4 |
| 4 | R-HSA-202733 | Cell surface interactions at the vascular wall | 5/61 | 137/11009 | 9,44E-04 | 8,47E-03 | 6,22E-03 | CD74/COL1A1/PPIA/COL1A2/FN1 | 5 |
| 4 | R-HSA-6802948 | Signaling by high-kinase activity BRAF mutants | 3/61 | 36/11009 | 1,01E-03 | 8,93E-03 | 6,56E-03 | ACTB/ACTG1/FN1 | 3 |
| 4 | R-HSA-2129379 | Molecules associated with elastic fibres | 3/61 | 37/11009 | 1,10E-03 | 9,20E-03 | 6,76E-03 | FN1/FBLN2/FBN1 | 3 |
| 4 | R-HSA-9735869 | SARS-CoV-1 modulates host translation machinery | 3/61 | 37/11009 | 1,10E-03 | 9,20E-03 | 6,76E-03 | RPS27L/EEF1A1/RPS12 | 3 |
| 4 | R-HSA-9678108 | SARS-CoV-1 Infection | 5/61 | 142/11009 | 1,11E-03 | 9,20E-03 | 6,76E-03 | PPIA/BST2/RPS27L/EEF1A1/RPS12 | 5 |
| 4 | R-HSA-6798695 | Neutrophil degranulation | 9/61 | 480/11009 | 1,21E-03 | 9,89E-03 | 7,27E-03 | S100A11/PPIA/BST2/HLA-B/B2M/CD63/HLA-C/EEF1A1/CST3 | 9 |
| 4 | R-HSA-76009 | Platelet Aggregation (Plug Formation) | 3/61 | 39/11009 | 1,28E-03 | 1,03E-02 | 7,54E-03 | COL1A1/COL1A2/FN1 | 3 |
| 4 | R-HSA-164940 | Nef mediated downregulation of MHC class I complex cell surface expression | 2/61 | 10/11009 | 1,32E-03 | 1,04E-02 | 7,62E-03 | HLA-A/B2M | 2 |
| 4 | R-HSA-5674135 | MAP2K and MAPK activation | 3/61 | 40/11009 | 1,38E-03 | 1,07E-02 | 7,84E-03 | ACTB/ACTG1/FN1 | 3 |
| 4 | R-HSA-196025 | Formation of annular gap junctions | 2/61 | 11/11009 | 1,61E-03 | 1,22E-02 | 8,96E-03 | ACTB/ACTG1 | 2 |
| 4 | R-HSA-9656223 | Signaling by RAF1 mutants | 3/61 | 43/11009 | 1,71E-03 | 1,27E-02 | 9,35E-03 | ACTB/ACTG1/FN1 | 3 |
| 4 | R-HSA-170834 | Signaling by TGF-beta Receptor Complex | 4/61 | 94/11009 | 1,79E-03 | 1,32E-02 | 9,66E-03 | STAT1/COL1A2/TGFBR2/FBN1 | 4 |
| 4 | R-HSA-1566948 | Elastic fibre formation | 3/61 | 44/11009 | 1,82E-03 | 1,32E-02 | 9,66E-03 | FN1/FBLN2/FBN1 | 3 |
| 4 | R-HSA-190873 | Gap junction degradation | 2/61 | 12/11009 | 1,92E-03 | 1,34E-02 | 9,83E-03 | ACTB/ACTG1 | 2 |
| 4 | R-HSA-430116 | GP1b-IX-V activation signalling | 2/61 | 12/11009 | 1,92E-03 | 1,34E-02 | 9,83E-03 | COL1A1/COL1A2 | 2 |
| 4 | R-HSA-2172127 | DAP12 interactions | 3/61 | 45/11009 | 1,95E-03 | 1,34E-02 | 9,83E-03 | HLA-B/B2M/HLA-C | 3 |
| 4 | R-HSA-190828 | Gap junction trafficking | 3/61 | 47/11009 | 2,21E-03 | 1,39E-02 | 1,02E-02 | ACTB/ACTG1/TUBA1B | 3 |
| 4 | R-HSA-437239 | Recycling pathway of L1 | 3/61 | 47/11009 | 2,21E-03 | 1,39E-02 | 1,02E-02 | ACTB/ACTG1/TUBA1B | 3 |
| 4 | R-HSA-6802946 | Signaling by moderate kinase activity BRAF mutants | 3/61 | 47/11009 | 2,21E-03 | 1,39E-02 | 1,02E-02 | ACTB/ACTG1/FN1 | 3 |
| 4 | R-HSA-6802949 | Signaling by RAS mutants | 3/61 | 47/11009 | 2,21E-03 | 1,39E-02 | 1,02E-02 | ACTB/ACTG1/FN1 | 3 |
| 4 | R-HSA-6802955 | Paradoxical activation of RAF signaling by kinase inactive BRAF | 3/61 | 47/11009 | 2,21E-03 | 1,39E-02 | 1,02E-02 | ACTB/ACTG1/FN1 | 3 |
| 4 | R-HSA-9649948 | Signaling downstream of RAS mutants | 3/61 | 47/11009 | 2,21E-03 | 1,39E-02 | 1,02E-02 | ACTB/ACTG1/FN1 | 3 |
| 4 | R-HSA-157858 | Gap junction trafficking and regulation | 3/61 | 49/11009 | 2,49E-03 | 1,54E-02 | 1,13E-02 | ACTB/ACTG1/TUBA1B | 3 |
| 4 | R-HSA-3928665 | EPH-ephrin mediated repulsion of cells | 3/61 | 51/11009 | 2,79E-03 | 1,71E-02 | 1,25E-02 | MMP2/ACTB/ACTG1 | 3 |
| 4 | R-HSA-75892 | Platelet Adhesion to exposed collagen | 2/61 | 15/11009 | 3,03E-03 | 1,82E-02 | 1,34E-02 | COL1A1/COL1A2 | 2 |
| 4 | R-HSA-449147 | Signaling by Interleukins | 8/61 | 473/11009 | 4,35E-03 | 2,56E-02 | 1,88E-02 | STAT1/MMP2/PPIA/VIM/COL1A2/ANXA1/FN1/PSMB4 | 8 |
| 4 | R-HSA-446353 | Cell-extracellular matrix interactions | 2/61 | 18/11009 | 4,36E-03 | 2,56E-02 | 1,88E-02 | ACTB/ACTG1 | 2 |
| 4 | R-HSA-202430 | Translocation of ZAP-70 to Immunological synapse | 2/61 | 19/11009 | 4,86E-03 | 2,81E-02 | 2,07E-02 | HLA-DRA/HLA-DPA1 | 2 |
| 4 | R-HSA-162909 | Host Interactions of HIV factors | 4/61 | 130/11009 | 5,76E-03 | 3,29E-02 | 2,42E-02 | PPIA/HLA-A/B2M/PSMB4 | 4 |
| 4 | R-HSA-164938 | Nef-mediates down modulation of cell surface receptors by recruiting them to clathrin adapters | 2/61 | 21/11009 | 5,93E-03 | 3,34E-02 | 2,46E-02 | HLA-A/B2M | 2 |
| 4 | R-HSA-6802952 | Signaling by BRAF and RAF1 fusions | 3/61 | 67/11009 | 6,03E-03 | 3,36E-02 | 2,47E-02 | ACTB/ACTG1/FN1 | 3 |
| 4 | R-HSA-202427 | Phosphorylation of CD3 and TCR zeta chains | 3/61 | 22/11009 | 6,50E-03 | 3,57E-02 | 2,62E-02 | HLA-DRA/HLA-DPA1 | 2 |
| 4 | R-HSA-140837 | Intrinsic Pathway of Fibrin Clot Formation | 2/61 | 23/11009 | 7,09E-03 | 3,80E-02 | 2,79E-02 | A2M/SERPING1 | 2 |
| 4 | R-HSA-389948 | PD-1 signaling | 3/61 | 23/11009 | 7,09E-03 | 3,80E-02 | 2,79E-02 | HLA-DRA/HLA-DPA1 | 2 |
| 4 | R-HSA-1445148 | Translocation of SLC2A4 (GLUT4) to the plasma membrane | 3/61 | 72/11009 | 7,36E-03 | 3,90E-02 | 2,87E-02 | ACTB/ACTG1/TUBA1B | 3 |
| 4 | R-HSA-1169408 | ISG15 antiviral mechanism | 3/61 | 73/11009 | 7,65E-03 | 4,00E-02 | 2,94E-02 | ISG15/STAT1/IFIT1 | 3 |
| 4 | R-HSA-9833482 | PKR-mediated signaling | 3/61 | 75/11009 | 8,24E-03 | 4,26E-02 | 3,13E-02 | ISG15/STAT1/TUBA1B | 3 |
| 4 | R-HSA-1169410 | Antiviral mechanism by IFN-stimulated genes | 4/61 | 149/11009 | 9,26E-03 | 4,74E-02 | 3,48E-02 | ISG15/STAT1/IFIT1/TUBA1B | 4 |
| 4 | R-HSA-9612973 | Autophagy | 4/61 | 151/11009 | 9,69E-03 | 4,90E-02 | 3,60E-02 | VIM/RNASE1/TUBA1B/EEF1A1 | 4 |
| 5 | R-HSA-2022090 | Assembly of collagen fibrils and other multimeric structures | 5/35 | 61/11009 | 1,26E-06 | 2,83E-04 | 2,32E-04 | COL6A3/COL15A1/COL4A2/COL3A1/PLEC | 5 |
| 5 | R-HSA-1474244 | Extracellular matrix organization | 8/35 | 300/11009 | 3,43E-06 | 2,83E-04 | 2,32E-04 | DCN/FBLN2/COL6A3/COL15A1/COL4A2/COL3A1/FBN1/PLEC | 8 |
| 5 | R-HSA-1474228 | Degradation of the extracellular matrix | 6/35 | 140/11009 | 4,55E-06 | 2,83E-04 | 2,32E-04 | DCN/COL6A3/COL15A1/COL4A2/COL3A1/FBN1 | 6 |
| 5 | R-HSA-390522 | Striated Muscle Contraction | 4/35 | 36/11009 | 4,69E-06 | 2,83E-04 | 2,32E-04 | TNNI2/TNNT3/TNNC2/VIM | 4 |
| 5 | R-HSA-264870 | Caspase-mediated cleavage of cytoskeletal proteins | 3/35 | 12/11009 | 6,35E-06 | 3,06E-04 | 2,51E-04 | GSN/PLEC/VIM | 3 |
| 5 | R-HSA-1474290 | Collagen formation | 5/35 | 90/11009 | 8,73E-06 | 3,51E-04 | 2,88E-04 | COL6A3/COL15A1/COL4A2/COL3A1/PLEC | 5 |
| 5 | R-HSA-8948216 | Collagen chain trimerization | 4/35 | 44/11009 | 1,06E-05 | 3,66E-04 | 3,00E-04 | COL6A3/COL15A1/COL4A2/COL3A1 | 4 |
| 5 | R-HSA-186797 | Signaling by PDGF | 4/35 | 58/11009 | 3,22E-05 | 9,69E-04 | 7,95E-04 | COL6A3/THBS4/COL4A2/COL3A1 | 4 |
| 5 | R-HSA-1442490 | Collagen degradation | 4/35 | 64/11009 | 4,75E-05 | 1,27E-03 | 1,04E-03 | COL6A3/COL15A1/COL4A2/COL3A1 | 4 |
| 5 | R-HSA-1650814 | Collagen biosynthesis and modifying enzymes | 4/35 | 67/11009 | 5,69E-05 | 1,37E-03 | 1,13E-03 | COL6A3/COL15A1/COL4A2/COL3A1 | 4 |
| 5 | R-HSA-3000178 | ECM proteoglycans | 4/35 | 76/11009 | 9,34E-05 | 2,05E-03 | 1,68E-03 | DCN/COL6A3/COL4A2/COL3A1 | 4 |
| 5 | R-HSA-216083 | Integrin cell surface interactions | 4/35 | 85/11009 | 1,44E-04 | 2,90E-03 | 2,38E-03 | COL6A3/COL4A2/COL3A1/FBN1 | 4 |
| 5 | R-HSA-111465 | Apoptotic cleavage of cellular proteins | 3/35 | 38/11009 | 2,30E-04 | 4,15E-03 | 3,41E-03 | GSN/PLEC/VIM | 3 |
| 5 | R-HSA-109581 | Apoptosis | 5/35 | 180/11009 | 2,41E-04 | 4,15E-03 | 3,41E-03 | GSN/C1QBP/PSMA2/PLEC/VIM | 5 |
| 5 | R-HSA-611105 | Respiratory electron transport | 4/35 | 103/11009 | 3,03E-04 | 4,68E-03 | 3,84E-03 | NDUFB5/NDUFB9/NDUFA7/UQCR10 | 4 |
| 5 | R-HSA-419037 | NCAM1 interactions | 3/35 | 42/11009 | 3,10E-04 | 4,68E-03 | 3,84E-03 | COL6A3/COL4A2/COL3A1 | 3 |
| 5 | R-HSA-397014 | Muscle contraction | 5/35 | 205/11009 | 4,39E-04 | 6,23E-03 | 5,11E-03 | TNNI2/TNNT3/TNNC2/ANXA1/VIM | 5 |
| 5 | R-HSA-5357801 | Programmed Cell Death | 5/35 | 213/11009 | 5,23E-04 | 7,01E-03 | 5,75E-03 | GSN/C1QBP/PSMA2/PLEC/VIM | 5 |
| 5 | R-HSA-75153 | Apoptotic execution phase | 3/35 | 52/11009 | 5,85E-04 | 7,42E-03 | 6,09E-03 | GSN/PLEC/VIM | 3 |
| 5 | R-HSA-163200 | Respiratory electron transport, ATP synthesis by chemiosmotic coupling, and heat production by uncoupling proteins. | 4/35 | 127/11009 | 6,70E-04 | 8,08E-03 | 6,63E-03 | NDUFB5/NDUFB9/NDUFA7/UQCR10 | 4 |
| 5 | R-HSA-6799198 | Complex I biogenesis | 3/35 | 57/11009 | 7,66E-04 | 8,79E-03 | 7,22E-03 | NDUFB5/NDUFB9/NDUFA7 | 3 |
| 5 | R-HSA-166658 | Complement cascade | 3/35 | 58/11009 | 8,06E-04 | 8,83E-03 | 7,25E-03 | CFD/C1R/SERPING1 | 3 |
| 5 | R-HSA-375165 | NCAM signaling for neurite out-growth | 3/35 | 63/11009 | 1,03E-03 | 1,07E-02 | 8,83E-03 | COL6A3/COL4A2/COL3A1 | 3 |
| 5 | R-HSA-3000480 | Scavenging by Class A Receptors | 2/35 | 19/11009 | 1,62E-03 | 1,63E-02 | 1,34E-02 | COL4A2/COL3A1 | 2 |
| 5 | R-HSA-1428517 | The citric acid (TCA) cycle and respiratory electron transport | 4/35 | 178/11009 | 2,34E-03 | 2,13E-02 | 1,75E-02 | NDUFB5/NDUFB9/NDUFA7/UQCR10 | 4 |
| 5 | R-HSA-140837 | Intrinsic Pathway of Fibrin Clot Formation | 2/35 | 23/11009 | 2,38E-03 | 2,13E-02 | 1,75E-02 | C1QBP/SERPING1 | 2 |
| 5 | R-HSA-166663 | Initial triggering of complement | 2/35 | 23/11009 | 2,38E-03 | 2,13E-02 | 1,75E-02 | CFD/C1R | 2 |
| 5 | R-HSA-8957275 | Post-translational protein phosphorylation | 3/35 | 108/11009 | 4,78E-03 | 4,12E-02 | 3,38E-02 | FSTL1/FBN1/CST3 | 3 |

**Supplemental Figure S1.** Selection of appropriate number of transcriptomics signatures K a priori. In general, the model perplexity (red curve) is decreasing depending on K, but an increasing perplexity is not problematic if the LDA shape parameter α < 1. The optimal K should theoretically minimize the perplexity in accordance with the number of rare transcriptomics signatures that could be unveiled (blue curve). After K=2 (irrelevant), the best models is found for K=5. See Stdeconvolve publication Miller et al. (2022) for more details.

**Supplemental Figure S2**. Spatialized transcriptomic pathological signatures are associated with myofibre injuries. Visium spots using 10X Genomics software Loupe Browser 7.0.1 in muscle cryosection from Acute (A JDM) (A) and Indolent (I JDM) (B) JDM, and healthy muscle (HM) (C). Each Visium spot was assigned to the molecular signature showing the maximal percentage after deconvolution. D-E: HE stained muscle cryosection from acute JDM showing ischemic injuries in acute JDM with microinfarcts, myosinolysis and perifascicular atrophy (D) and from indolent slowly progressive JDM showing chronic moderate myopathic changes with myofiber size variation and internalized nuclei (E).

**Supplemental Figure S3.** Ten first enriched Reactome pathways. Queries were the up regulated genes from marker genes list (log2fc > 1) for signature 2 unveiled by de novo deconvolution.
